# Hb-EGF directs systemic muscle repair

**DOI:** 10.1101/2025.06.04.657846

**Authors:** Haley C. Dean, Vishnu M. Saraswathy, Amulya Saini, Tiffany Ou, Jennifer McAdow, Amruta Tendolkar, Mayssa H. Mokalled, A.N. Johnson

**Affiliations:** Department of Developmental Biology, Washington University in St. Louis, St Louis, MO 63108

**Keywords:** regenerative capacity, muscle regeneration, systemic muscle injury, Hb-EGF, zebrafish

## Abstract

Regenerative capacity varies between tissues, species, and stages of the life cycle. What is less appreciated is that regenerative capacity also varies with the magnitude of the injury, even within a single tissue. Vertebrate skeletal muscle efficiently regenerates following minor injuries; however, extensive injuries may result in incomplete repair, which can be debilitating. To understand if small- and large-scale muscle injuries activate distinct regenerative programs, we developed a systemic muscle injury model in zebrafish. Transcriptomic analysis of muscle and non-muscle tissues revealed that systemic and local muscle injuries elicit distinct molecular responses, both quantitatively and qualitatively. Systemic muscle injury activated the expression of Heparin binding epidermal-like growth factor (Hb-EGF) in the epidermis, and Hb-EGF is necessary for systemic muscle repair. Conversely, local muscle injury did not induce Hb-EGF expression and Hb-EGF was not required for local muscle repair. These studies suggest that large- and small-scale muscle injuries activate different regenerative programs, resulting in either systemic or local repair.

## Introduction

Regenerative capacity varies significantly across species and developmental stages. Planaria and hydra can regenerate entire body axes (Morgan, 1898; Morgan, 1901; Gierer et al., 1972; Zammit, 2017), amphibians can regenerate entire limbs and organs (Fraisse, 1885; Harrison, 1921; Tanaka, 2003; Roddy et al., 2008), while mammalian regenerative capacity in the adult is limited to just a few tissues including the skin, liver, bone, and skeletal muscle. As mammals age regenerative capacity is lost; for example fracture healing and muscle repair are slowed in the aging population, which contributes to a significant burden on the healthcare system and impacts the quality of life for older adults (Saul and Khosla, 2022; Grima-Terrén et al., 2024; Nguyen et al., 2025). While extensive comparative studies have revealed mechanisms that confer differential regenerative capacity, it remains unclear if the magnitude of the injury itself activates distinct regenerative programs in tissues and organisms with high regenerative capacity.

Transection injury models have provided some insights into the organismal response to injuries of different magnitudes. Planarians regenerate a new head after amputation, but do not duplicate the spared tissues or create additional body axes (Morgan, 1904b; Morgan, 1904a). This foundational observation argues planarians have the capacity to recognize the extent of tissue loss and then elicit the appropriate regenerative response. Planarians repair large-scale injuries by activating a “missing tissue response” that accelerates regeneration to replace the lost tissue (Tewari et al., 2018). The rate of regenerative growth in response to forelimb amputation in newts is also dependent on the magnitude of the injury. The regenerate produced after forelimb amputation proximal to the body grows faster than regenerates produced by distal amputations (Iten and Bryant, 1973). Similarly, regeneration rate of zebrafish caudal fin rays following amputation correlates with the length of fin ray that is being replaced (Uemoto et al., 2020). Modulating the regeneration of individual rays reproduces a “forked” caudal fin when all rays are amputated along a single plane. These studies suggest the magnitude of tissue injury in highly regenerative organisms is sensed by a rheostat mechanism that governs the regeneration growth rate to appropriately compensate for the lost tissue.

Even though differential regeneration rates can compensate for the magnitude of tissue loss in certain contexts, some regenerative organs are limited in the volume of tissue that can be replaced following injury. The adult zebrafish heart for example can regenerate a ventricular resection, but only if less than 20% of the ventricular volume is lost (Poss et al., 2002). Similarly, neonatal mice can regenerate a ventricular amputation that removes up to 15% of the ventricular mass (Porrello et al., 2011). Skeletal muscle is also highly regenerative, but unlike heart regeneration which depends on the dedifferentiation and proliferation of spared cardiomyocytes (Kikuchi et al., 2010), skeletal muscle repair depends on dedicated populations of muscle stem cells (MuSCs). Muscle injury activates MuSCs, which undergo asymmetric divisions to replenish the stem cell pool as well as transient amplification and differentiation to generate myoblast progenitors dedicated to muscle repair (Troy et al., 2012; Pawlikowski et al., 2017; Feige et al., 2018). Mature myoblasts can fuse with and repair surviving myofibers or fuse with each other to replace damaged myofibers with new myofibers (Demonbreun et al., 2015). MuSC-dependent regeneration can repair small-scale injuries, including targeted mechanical ablation in zebrafish or toxin-based injuries of a single muscle in mammals (Gurevich et al., 2016; Hardy et al., 2016; Ratnayake et al., 2021). However, large-scale muscle loss in mammals that exceeds 20% of muscle mass results in incomplete muscle repair and often long-term disability (Sicherer et al., 2020). The striking contrast in regenerative capacity between amputated limbs and resected striated muscles argues accelerating regenerative growth rates alone is not sufficient to repair all large-scale injuries.

One unexplored mechanism that could explain differential regenerative capacity is that independent regenerative programs direct organ repair in response to small- and large-scale injuries. We developed an inducible skeletal muscle injury model in zebrafish that damages all skeletal muscles simultaneously. Injured larvae showed a dramatic loss of motor function two days after systemic muscle injury and strikingly, injured animals regained full motor function ten days later. A model of local muscle injury and repair had been established for zebrafish larvae (Gurevich et al., 2016; Ratnayake et al., 2021), and we used single-cell RNA sequencing (scSeq) of muscle and non-muscle tissues to compare the molecular responses to systemic and local muscle injuries. Our analysis identified quantitative differences in nearly 90 percent of the signaling pathways between systemic or local muscle injuries. Computational analysis suggested Heparin binding epidermal like growth factor (Hb-EGF) signals between the epidermis and muscle stem cells in response to systemic but not local muscle injury. Our subsequent mutational analysis showed Hb-EGF is required for systemic repair but is dispensable for local repair. These studies support a model in which large- and small-scale muscle injuries activate distinct regenerative programs that direct systemic or local repair.

## Results

### An inducible injury model to investigate systemic muscle repair

To understand if systemic and local muscle injuries activate distinct regenerative programs, we developed an inducible systemic muscle injury model. The nitroreductase (NTR)- metronidazole (MTZ) system uses tissue-specific NTR expression to induce targeted injuries in zebrafish (Curado et al., 2008). NTR is innocuous under physiological conditions in the absence of MTZ. However, in the presence of MTZ, NTR converts the MTZ pro-drug into a DNA crosslinking protein that activates the apoptotic machinery. We used the strong *actin alpha cardiac muscle1b* (*actc1b*) skeletal muscle promoter to express NTR and mCherry in mature myofibers [hereafter Tg(*actc1b:*NTR,2A,mCherry)]. This construct did not activate mCherry expression in the heart, suggesting NTR-induced injury would be skeletal muscle specific. Larval muscles injured at 4 days post fertilization (dpf) with local needle-stick injury show efficient muscle repair by 6dpi (Gurevich et al., 2016). We treated 4dpf Tg(*actc1b:*NTR,2A,mCherry) larvae with a range of MTZ concentrations and incubation times and found 12hr treatment of 5mM MTZ drastically reduced swim function at 2dpi, but strikingly the injured larvae regained full swim function by 12dpi (Fig. 1A,B). Zebrafish larvae are therefore able to repair systemic muscle injuries over a time course similar to that of local muscle injuries.

**Figure 1.**
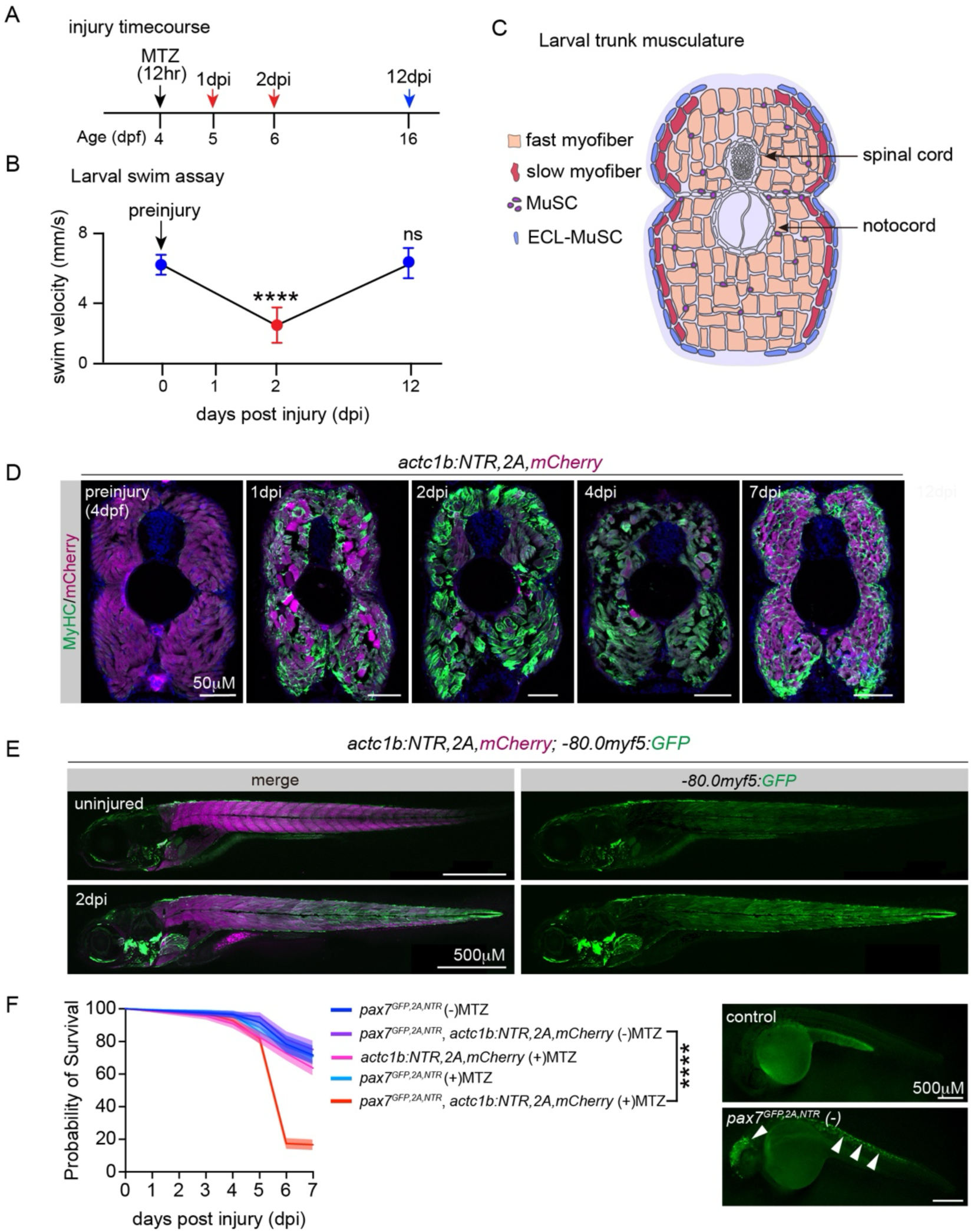
An inducible model of systemic skeletal muscle injury. **A.** Experimental design to induce muscle injury and assess regeneration**. B.** Larval swim function assay. Swim function of injured Tg(*actc1b*:NTR,2A,mCherry) larvae was drastically impaired at 2dpi but was restored by 12dpi. n≥19 per cohort. **C.** Diagram of a transverse section through the larval trunk musculature, showing the cellular composition of the myotome. Dorsal is oriented toward the top. **D.** Regeneration assay. Transverse sections of the trunk musculature of Tg(*actc1b*:NTR,2A,mCherry) larvae labelled for mCherry (red) and MyHC (green). At 1dpi MyHC begins to aggregate in myofibers throughout the trunk musculature. MyHC aggregates are resolved ventrally by 7dpi and MyHC is enriched at the myofiber cortex similar to uninjured controls. **E.** Live images of larvae double transgenic for Tg(*actc1b*:NTR,2A,mCherry),(*-80.0myf5*:GFP) treated with MTZ for 12 hours at 4dpf. Injured larvae showed increased *myf5*:GFP expression (green) at 2dpi. **F.** Survival curve. *pax7a^GFP,2A,NTR2.0^*Tg(*actc1b*:NTR,2A,mCherry) larvae treated with MTZ showed significantly reduced survival. Live images show control and *pax7a^GFP,2A,NTR2.0^* embryos at 26hpi. Arrows denote GFP+ cells. Significance was determined by unpaired students t-test (**B**) and Log-rank test (**F**). Error bars represent SEM. **** (p<0.0001).

At the cellular level, Myosin Heavy Chain (MyHC) aggregated in myofibers throughout the trunk musculature at 2dpi which likely reflects sarcomere disassembly (Fig. 1C,D). By 7dpi MyHC aggregates had been cleared from the trunk musculature and MyHC instead localized to the myofiber cortex (Fig. 1D). Regenerating myofibers in mammals are easily distinguished by the presence of centralized myonuclei, but we did not observe centralized nuclei in myofibers following systemic injury. mCherry expression from the Tg(*actc1b:*NTR,2A,mCherry) transgene was reduced in injured myofibers at 2dpi but then restored by 7dpi (Fig. 1D). Myf5 is expressed in activated MuSCs and proliferating muscle progenitors during muscle regeneration (Gurevich et al., 2016). Injured larvae transgenic for Tg(*actc1b:*NTR,2A,mCherry) and Tg(*myf5*:GFP) activated GFP expression at 2dpi, suggesting systemic repair is a MuSC-based process (Fig. 1E). To functionally test if MuSCs are required for systemic repair, we knocked an NTR,2A,GFP cassette into the *pax7a* locus to generate *pax7a^GFP,2A,NTR2.0^* fish. Pax7 is expressed in quiescent and activated MuSCs and *pax7a^GFP,2A,NTR2.0^*larvae treated with MTZ are expected to have fewer MuSCs than control larvae. We treated Tg(*actc1b:*NTR,2A,mCherry) and *pax7a^GFP,2A,NTR2.0^*Tg(*actc1b:*NTR,2A,mCherry) larvae with MTZ and found *pax7a* knock-in larvae had significantly reduced viability compared to controls (Fig. 1F). These data argue systemic muscle injury reduces motor function by disrupting the myofiber contractile machinery and that systemic repair is accomplished through a MuSC-dependent process.

### Systemic regeneration repairs surviving myofibers

*slow myosin heavy chain 1* (*smyhc1*) is expressed in newly created fast myofibers after local muscle injury in larvae (Gurevich et al., 2016), arguing zebrafish can repair muscle through multiple cellular mechanisms. To understand whether systemic repair produces *de novo* myofibers, we generated a *smyhc1^sfgfp^* knock-in line (Fig. S1A). Uninjured *smyhc1^sfgfp^* larvae expressed GFP in the superficial slow myofibers of the truck musculature but not in the deep fast myofibers (Fig. S1A,B). GFP expression in slow muscles was drastically altered by systemic injury, but surprisingly GFP+ myofibers were not detected in the fast myotome during systemic repair (Fig. S1A,B). Systemic repair therefore does not create new myofibers.

To further investigate myofiber fate after systemic injury, we generated Tg*(actc1b:*CreER^T2^) transgenic fish that express Tamoxifen (TMX)-dependent CreER^T2^ in myofibers for lineage tracing experiments (Fig. 2A). We treated larvae that were triple transgenic for Tg(*actc1b:*NTR), Tg*(actc1b:*CreER^T2^), and the lineage trace reporter Tg(*ubb3*:RFP.loxP.STOP.loxP.GFP) at 2dpf with TMX for 24hr to induce recombination of the lineage reporter. GFP expression was broadly induced throughout the trunk musculature 48 hours after TMX treatment (Fig. 2B). The prediction for the lineage trace experiment was that damaged myofibers that are repaired will remain GFP+ while damaged myofibers that are replaced with new myofibers will be GFP negative (Fig. 2C). We injured TMX-treated triple transgenic larvae with MTZ for 24hr and live imaged GFP fluorescence for 24hr. Injured myofibers maintained GFP expression during 24hr of live-imaging, during which there were no indications of myofiber death or clearance (Fig. S2A). Syncytial myofibers in flies and newts can cellularize to produce reparative, mononucleate myoblasts that proliferate and generate new myofibers (Frasch, 2016), but we saw no evidence of myofiber cellularization during live-imaging. We extended the lineage trace analysis out to 7dpi using fixed tissue, and found the number of GFP+ myofibers at 2-, 4-, and 7dpi were comparable between injured and uninjured larvae (Fig. 2D). The absence of *smyhc1^sfgfp^*expression in fast myofibers during systemic repair and the persistence of GFP expression in lineage traced myofibers after injury argue the primary cellular mechanism of systemic muscle repair is fusion-based repair of surviving myofibers.

**Figure 2.**
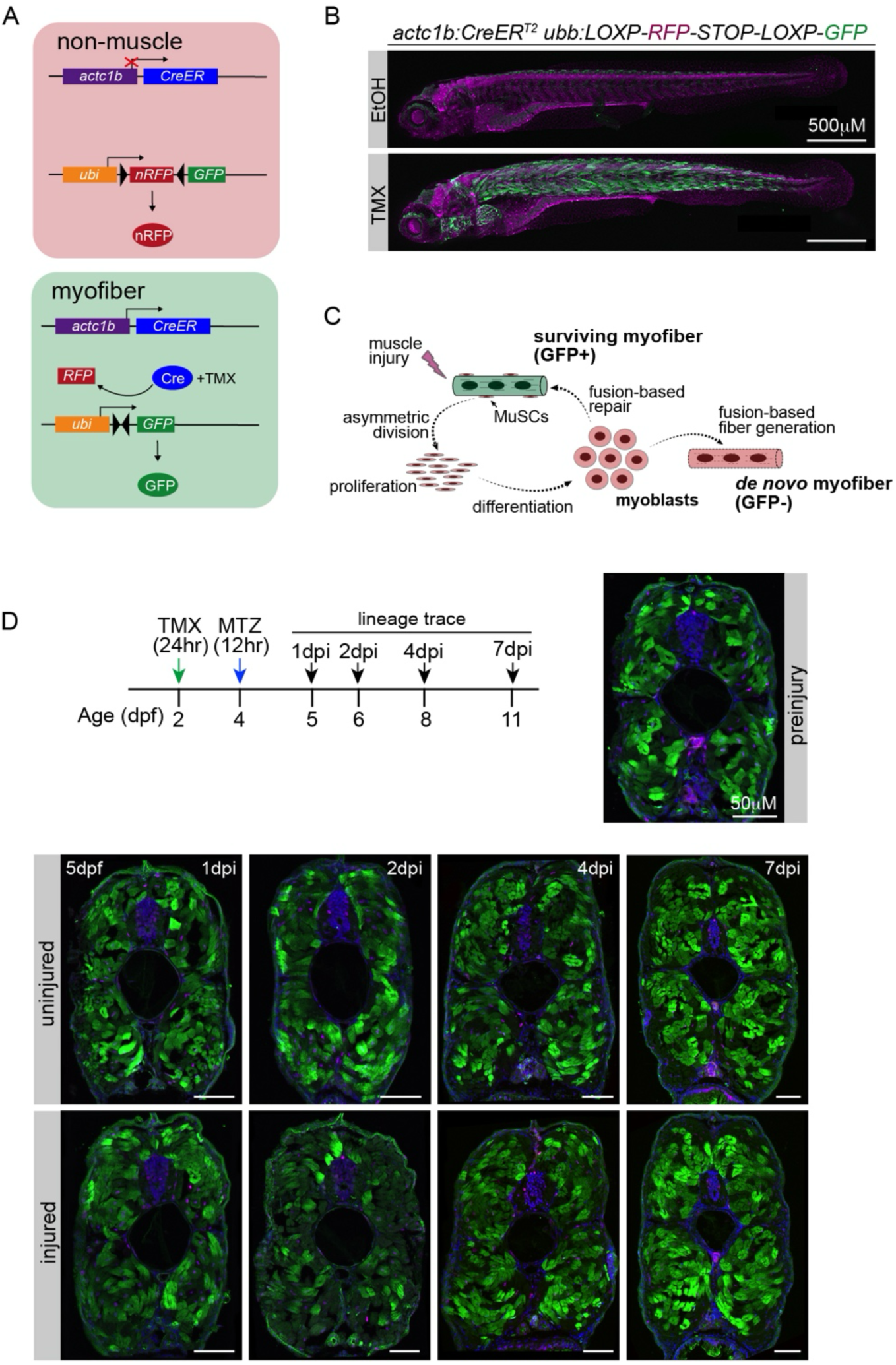
Injured myofibers are repaired after systemic injury. **A.** Lineage tracing strategy. Upon treatment with tamoxifen (TMX), *actc1b*:CreER^T2^ induces recombination in the target cassette, which activates GFP expression in myofibers. **B.** Live images of 5dpf larvae double transgenic for Tg(*actc1b*:CreER^T2^),(*ubb*:LOXP-RFP-STOP-LOXP-GFP) 24hrs after TMX treatment. TMX-treated larvae showed GFP+ myofibers (TMX) but vehicle treated larvae (EtOH) did not activate GFP expression. RFP protein is highly stable and continues to fluoresce after recombination. **C.** Schematic of muscle regeneration in lineage traced myofibers. After muscle injury, surviving myofibers that are repaired remain GFP+ while *de novo* myofibers are GFP-. **D.** Experimental strategy for lineage tracing over 7d. 2dpf larvae triple transgenic for Tg (*actc1b*:NTR), (*actc1b*:CreER^T2^),(*ubb*:LOXP-RFP-STOP-LOXP-GFP) were treated with TMX for 24 hours. Injury was MTZ-induced at 4dpf followed by sample collection at 2-, 4- and 7 dpi. Transverse sections of lineage traced larvae revealed injured and uninjured fish had a comparable number of GFP+ and GFP-myofibers in the trunk musculature over the experimental time course. A vast majority of myofibers survive systemic injury and are repaired. See **Fig. S2B** for lineage trace quantification.

### Systemic and local injuries activate distinct signaling pathways

Systemic and local muscle repair likely involve distinct regenerative programs. To evaluate the molecular responses to systemic and local injuries, we performed single cell RNA sequencing (scSeq) of uninjured and MTZ-injured larvae at 1-, 2-, and 4-dpi, and scSeq of locally injured (needlestick) larvae at 2dpi. Cell suspensions of dissociated trunks from multiple individuals were pooled for each cohort, and isolated cells were sequenced using 10X genomics platform (3’ v3.1 chemistry). Sequence reads were aligned to zebrafish genome GRCz11, and cells were filtered using the Decontx and DoubletFinder packages to exclude cells that include doublets or high quantities of ambient and mitochondrial RNA (see Methods for a complete description of scSeq bioinformatic analyses). A total of 67,554 cells were used for subsequent analyses.

Unsupervised clustering identified 32 cell populations in the larval trunk (Fig. 3A). We used the differentially expressed genes (DEGs) among cell populations from a zebrafish embryo and larva scSeq atlas(Sur et al., 2023) to generate a matrix that assigned the 32 clusters to one of 14 tissues (Fig. S3A, Table S1). Six of the clusters were assigned to multiple tissues, which we resolved by comparing only the identifier genes from the scSeq atlas to each tissue type (Fig. S3B). Clusters were then assigned subidentities using cell type specific markers from the scSeq atlas (Fig. 3B). In all, five clusters of muscle cells were identified (MuSCs, differentiating MuSCs, slow myofibers, and two clusters of fast myofibers; Fig. S3C), along with other cell types expected to be present in the larval trunk (cartilage, tendons, neurons, periderm, vasculature, and monocytes). Subclustering the muscle cell populations uncovered seven MuSC clusters, which showed differential expression of the MuSC markers *en1* and *pax7a* (Fig. 3C,D). The muscle subclusters also contained differentiating muscle progenitors, that expressed *myod* and *myog*, and terminally differentiated myofibers that expressed *actc1b* (Fig. S3D). Comparing the subclusters to single cell and single nucleus studies of regenerating mouse muscles (McKellar et al., 2021), revealed two populations of activated MuSCs (Fig. 3C). Notably, the number of activated MuSCs was greater in larvae with systemic injuries than in uninjured larvae or in larvae with local injuries (Fig. 3F). These studies argue systemic injuries activate more MuSCs to complete a large-scale repair than local injuries.

**Figure 3.**
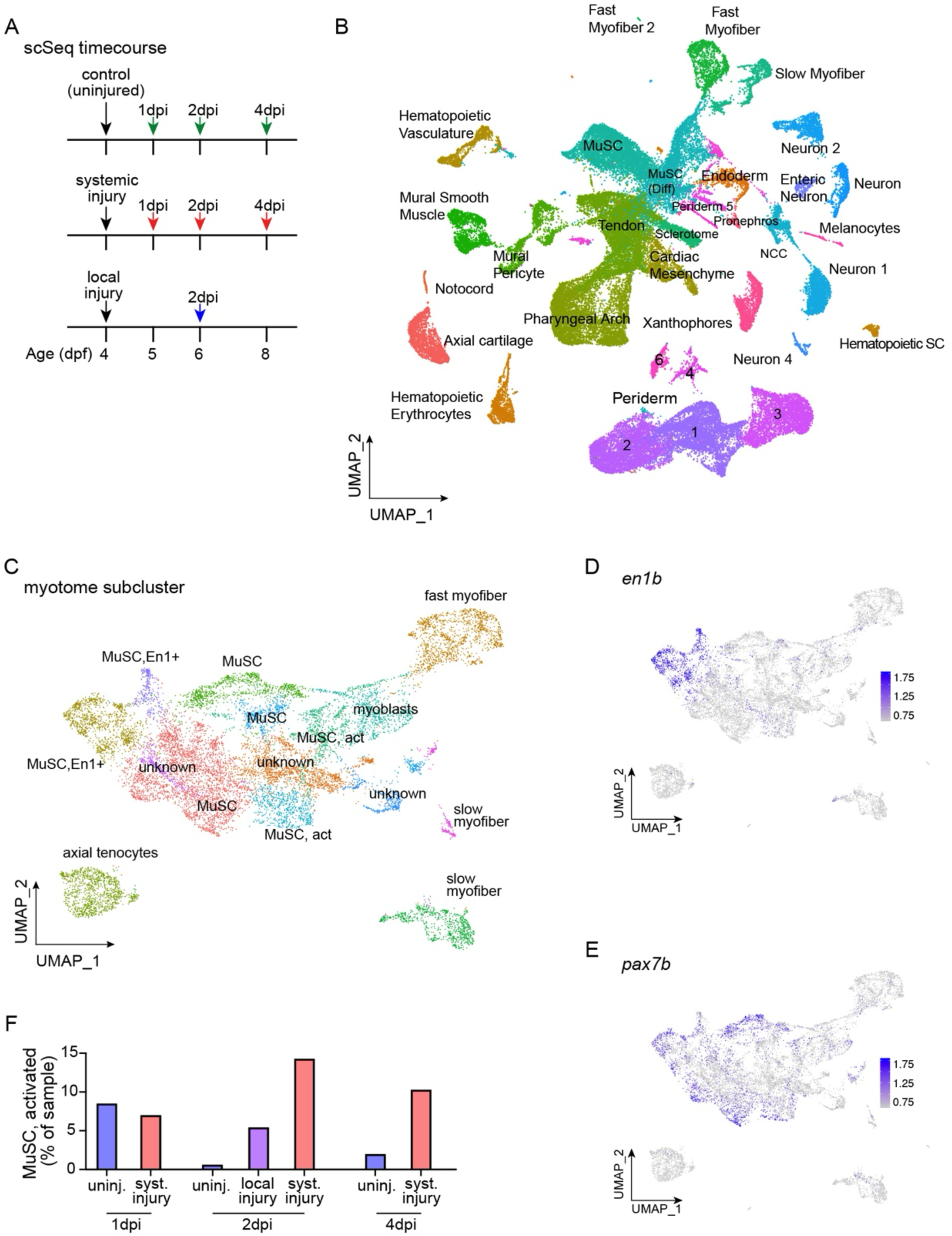
scSeq of larvae with systemic and local muscle injuries. **A.** Experimental design for single cell sequencing (scSeq). Seven cohorts were collected for scSeq. **B.** Merged UMAP representation of the complete data set, representing 67,554 cells. A total of 31 clusters were identified. **C.** UMAP plot showing 15 muscle subclusters, including multiple population of MuSCs. The precise identities of three clusters remain unknown. **D-E.** Feature plots showing the distributions of canonical MuSC markers *en1b* (**D**) and *pax7b* (**E**) among the muscle subclusters. **F.** Distribution of activated MuSCs during muscle repair. Cell proportions were normalized to the total number of cells at each time point. Systemic injury produced more MuSCs than local injury.

To understand if systemic and local injuries induce distinct signaling responses, we used the CellChat program to quantify intercellular communication networks based on the expression of ligands, receptors, and pathway modifiers (Jin et al., 2021; Jin et al., 2025). Zebrafish genes were first converted to human orthologues, as we previously described (Saraswathy et al., 2024), and then CellChat was used to calculate the communication probabilities of individual signal transduction pathways in outgoing (signaling) and incoming (signal receiving) cells. The cumulative probabilities of outgoing and incoming signals were then integrated across all pathways, which revealed the cumulative interaction strength was higher in response to systemic injury than local injury (Fig. 4A). The cumulative outgoing and incoming signaling strength was also higher in response to systemic injury (Fig. 4B). Cumulative signaling strength can also be calculated for individual clusters. Four clusters (periderm1, MuSCs, neuron4, and sclerotome) showed dramatically higher incoming and outgoing signaling strength in response to systemic injury than local injury (Fig. S4A).

**Figure 4.**
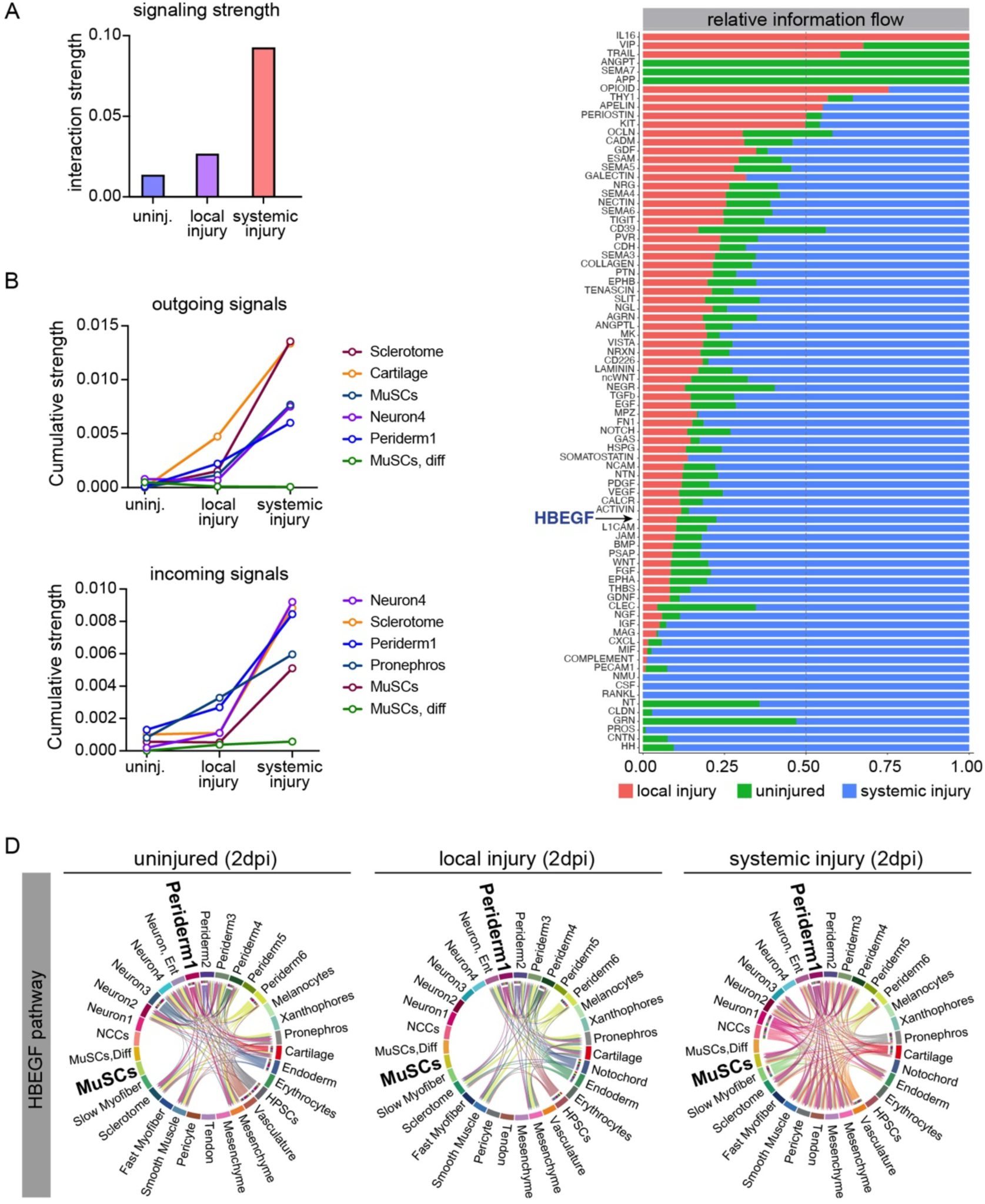
Systemic injury activates unique cell-cell communication networks. **A.** Relative interaction strengths in uninjured (uninj.), locally injured, and systemically injured samples at 2dpi. **B.** Cumulative strengths of signaling pathways outgoing from or received by the five strongest cell types at 2dpi. The “MuSC, diff” (differentiated MuSCs) population is shown as a low-strength outlier. **C.** Information flow plot showing relative signaling flow among the three samples at 2dpi. Most signaling pathways showed greater information flow in response to systemic injury, including Heparin binding epidermal-like growth factor (Hb-EGF). D. Chord diagrams showing Hb-EGF intercellular communication in response to injury at 2dpi. Note only systemic injury induces Hb-EGF signaling to MuSCs.

Next, we compared the information flow for each signaling pathway between uninjured, local, and systemic injuries (Fig. 4C). Information flow is the sum of communication probabilities across clusters and conditions for an individual pathway. We found that several signaling pathways showed information flow exclusively under uninjured (ANGPT, SEMA7, and APP), locally injured (IL16), or systemically injured (CSF, NMU, RANKL) conditions, but a majority of the signaling pathways showed dynamic information flow across conditions (Fig. 4C). In addition, information flow for 62 of the remaining 74 signaling pathways was higher in systemic injury than local injury or injured samples. These analyses suggest systemic and local injuries induce distinct signaling responses that activate qualitatively different regenerative programs.

### Systemic injuries activate signaling pathways outside of the MuSC niche

To determine which signaling pathways might control unique systemic repair programs, we looked for pathways with high information flow in systemic injury and that either signaled to MuSCs only in response to systemic injury or that signaled out of MuSCs only in response to systemic injury. Three signaling pathways met these criteria including the Epidermal growth factor (EGF), Macrophage migration inhibitory factor (MIF), and Nerve growth Factor (NGF). EGF and MIF are MuSC incoming signals and NGF is a MuSC outgoing signal (Fig. S4B). To understand which EGF ligand might be signaling to MuSCs after systemic injury, we repeated the CellChat analysis using individual ligands and found Heparin binding epidermal growth factor (Hb-EGF) signaling recapitulated the outputs observed using all EGF ligands (Figs. 4D, S4B). Periderm1 and MuSCs had some of the highest cumulative interaction strengths among the clusters (Fig. S4A), and Hb-EGF showed the strongest signaling from periderm1 to MuSCs (Fig. 4D). Hb-EGF was thus a strong candidate for a unique signaling response that is activated by systemic but not local injury.

We had previously developed a high throughput pipeline to validate scSeq candidates in vivo using expression analysis and mutagenesis (Saraswathy et al., 2024). Hybridization Chain Reaction (HCR)-fluorescent *in situ* hybridization (HCR-FISH) is first used to evaluate transcript expression in response to injury. The requirement of injury-induced transcripts during regeneration is then tested using ribonucleotide (RNP)-mediated CRISPR genome editing. The major advantage of this validation protocol is that the mutagenic efficiency of RNP CRISPR technology is sufficient to screen F_0_ animals. The zebrafish genome encodes two *hbegf* transcripts, *hbegfa* and *hbegfb*, but scSeq analysis showed only *hbegfa* was induced in the periderm at 1-, 2-, and 4-days after systemic injury (Fig. 5A). HCR-FISH in Tg(*myf5*:GFP) larvae confirmed *hbegfa* is induced by systemic injury and is expressed superficial to the external cell layer at a location consistent with the periderm (Fig. 5B). Importantly, *hbegfa* was not expressed in Myf5+ MuSCs or muscle progenitors in response to systemic injury. We extended our HCR analysis to compare *hbegfa* expression after systemic and local injury. Local injury is consistently performed at the same position along the rostral-caudal axis. We used this rostral-caudal position to quantify *hbegfa* expression at the local injury site and at an analogous position in larvae recovering from systemic injury. Incredibly, *hbegfa* was not induced at the local injury site but showed trending enrichment in the analogous position in response to systemic injury (Fig. 5C). These expression studies argue *hbegfa* is specifically induced outside the MuSC niche in response to systemic injury.

**Figure 5.**
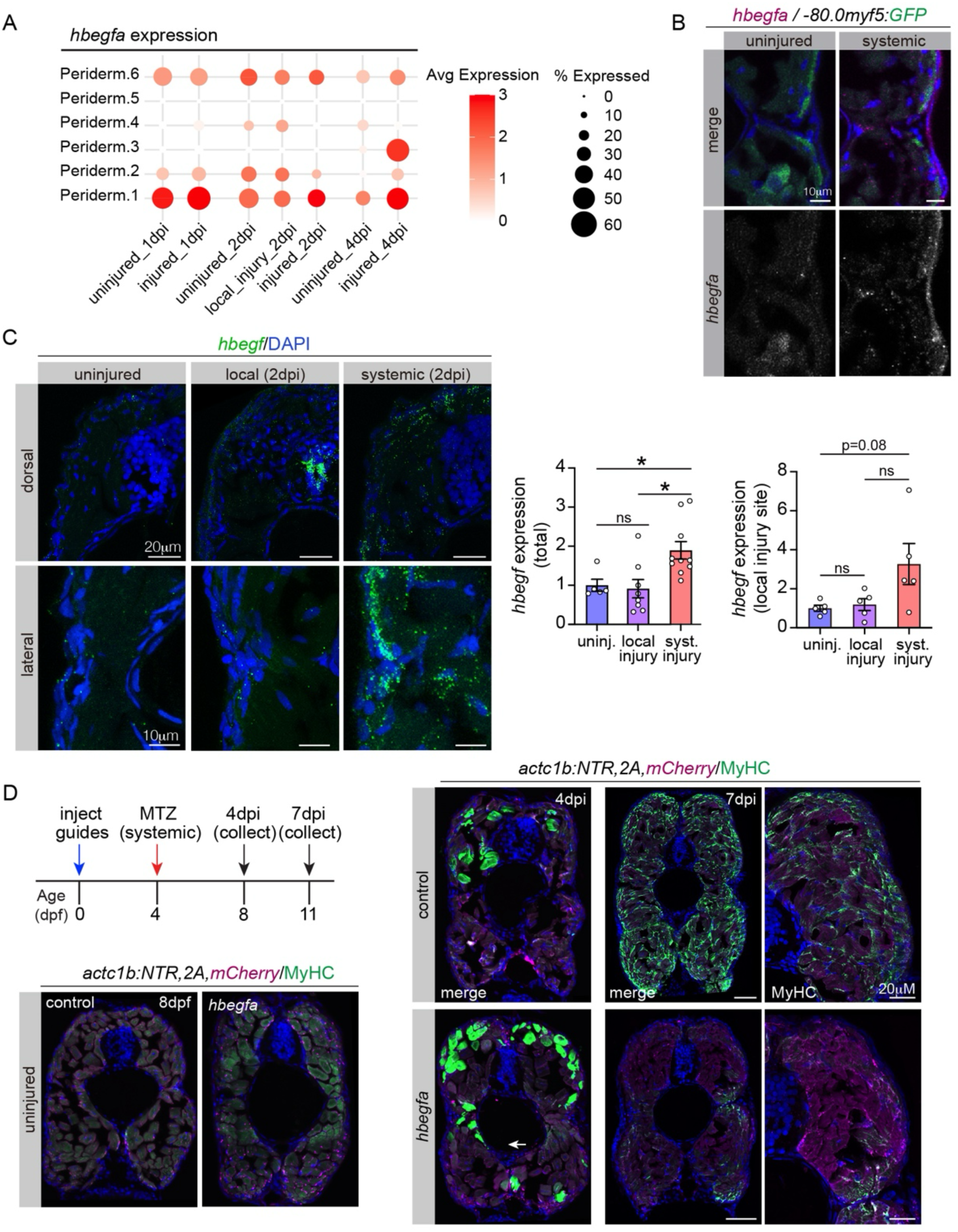
Systemic repair requires *hbegfa*. **A.** Dot plot of *hbegfa* expression in the periderm from the complete data set. Dot plot shows relative expression in each cell type. Dot colors and diameters represent average gene expression and the percent of cells expressing *hbegfa*. **B.** *hbegfa* HCR in larvae double transgenic for Tg(*actc1b*:NTR,2A,mCherry)(*-80.0myf5*:GFP) at 2dpi. *hbegfa* is induced superficial to Myf5+ MuSCs in response to systemic injury. **C.** *hbegfa* HCR in Tg(*actc1b*:NTR,2A,mCherry) larvae at 2dpi. *hbegfa* expression is induced in response to systemic but not local injury. Quantification was performed on complete transverse sections (left) or at the site of needlestick injury (right). Data points represent average expression values from individual larva. Error bars represent SEM. Unpaired t-test. (*) p<0.05. **D.** Experimental design for CRISPR-mediated deletion of *hbegfa*. Tg(*actc1b*:NTR,2A,mCherry) embryos were injected at the one cell stage, injured at 4dpf, and collected for histology. Transverse sections of control and *hbegfa* crispant larvae at 4- and 7dpi, labelled for MyHC (green) and mCherry (red). MyHC expression was reduced at 7dpi in *hbegfa* crispants. Uninjected siblings were used as controls.

### *hbegfa* is required for systemic repair

To understand if Hb-EGF is required for systemic repair, we used RNP CRISPR to delete the entire *hbegfa* coding region. The editing efficiency of each guide was greater than 50%, so we injured F_0_ *hbegfa* “crispants” double transgenic for Tg(*actc1b:*NTR,2A,mCherry) and *smyhc1^sfgfp^*at 4dpf and then live imaged slow muscle at 7dpi. Slow myofiber regeneration was significantly disrupted in *hbegfa* crispants compared to controls, with phenotypes that included missing and morphologically aberrant slow myofibers (Fig. S5A,B). Histological analysis of fast myofiber regeneration in Tg(*actc1b:*NTR,2A,mCherry) showed MyHC expression was dramatically reduced in 7dpi *hbegfa* crispants compared controls (Fig. 5D). In addition, MyHC did not localize to the myofiber cortex in *hbegfa* crispants, suggesting myofiber repair was impaired (Fig. 5D). Notably, uninjured *hbegfa* crispants showed normal muscle morphology at 11dpf, arguing the muscle phenotypes in injured *hbegfa* crispants result from defects in systemic repair rather than issues in muscle development or physiology. Hb-EGF is therefore required for systemic repair of fast and slow myofibers.

Our expression studies showed Hb-EGF expression is induced by systemic injury but not by local injury, suggesting Hb-EGF is dispensable for local muscle repair. We performed local injuries on *hbegfa* crispants and control injected larvae transgenic for *smyhc1^sfgfp^* and live imaged slow myofibers at 7dpi. Slow myofiber regeneration was comparable between *hbegfa* crispants and injured controls (Fig. S5A,B). These studies are consistent with the model in which Hb-EGF activates a systemic muscle repair program that is distinct from the local muscle repair program.

### Hb-EGF promotes muscle stem cell activation

*pax7b* and *en1b* are expressed in partially overlapping populations of MuSCs (Fig. 3D,E). We used En1 and Pax7 antibodies to visualize MuSC activity in response to systemic injury. En1 is expressed in spatially segregated MuSCs, known as the external cell layer (ECL), which are located superficial to slow myofibers and are thus spatially and molecularly distinct from interstitial MuSCs that do not express En1 (Figs. S6A)(Keenan and Currie, 2019). *hbegfa* crispants transgenic for Tg(*actc1b:*NTR,2A,mCherry) showed a significant reduction in the number of Pax7+ cells at 1- and 2-dpi compared to controls, but the number of En1+ cells was comparable between the two groups (Figs. 6A,B S6A). Importantly, the number of Pax7+ cells was not significantly different between *hbegfa* crispants and control larvae prior to injury, suggesting Hb-EGF is required for MuSC proliferation after injury (Fig. 6B).

**Figure 6.**
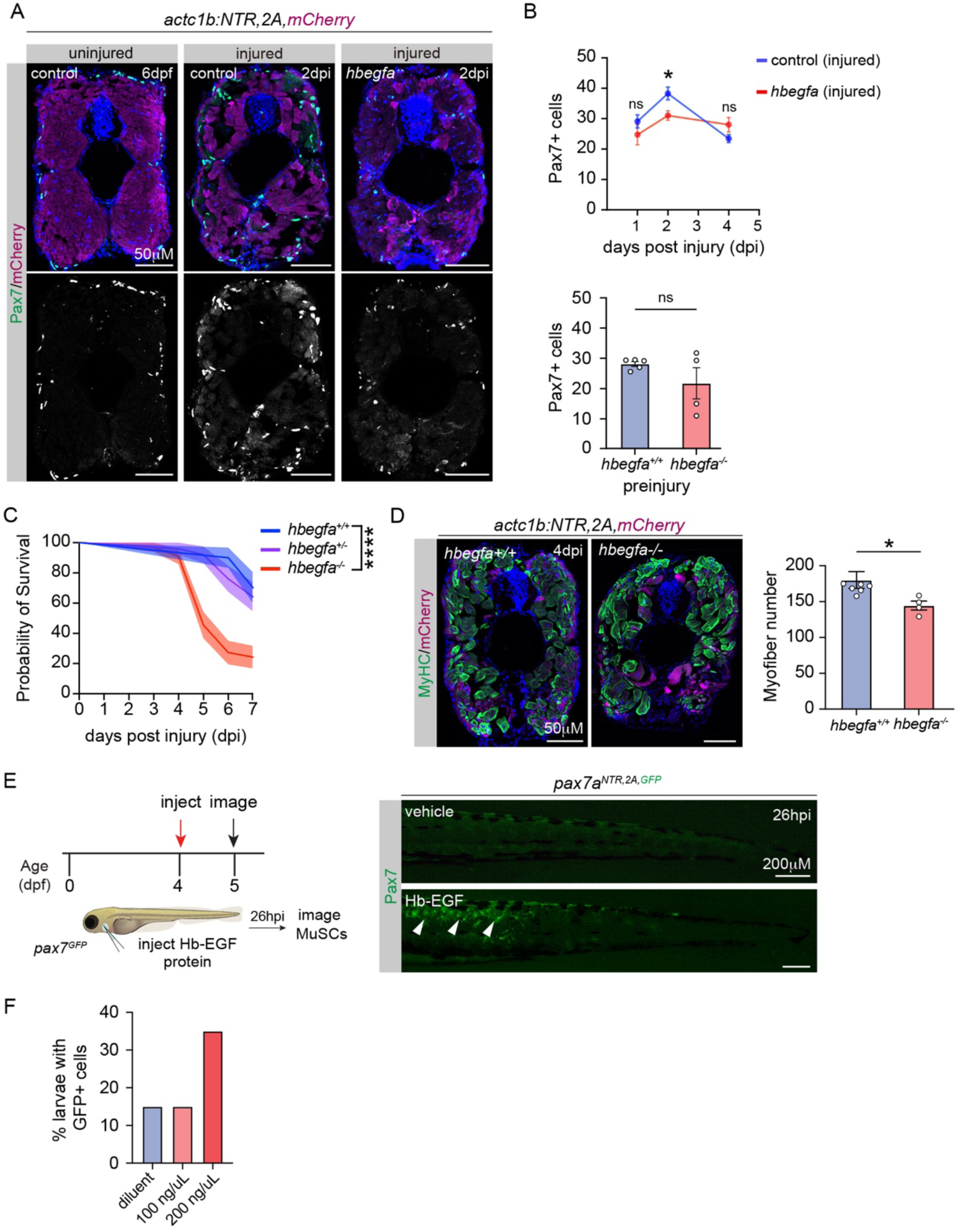
Hb-EGF regulates the MuSC response to systemic injury. **A.** Transverse sections of Tg(*actc1b*:NTR,2A,mCherry) larvae at 6dpf (uninjured) or 2dpi (injured), labelled for Pax7 (green) and mCherry (red). *hbegfa* crispants were injected at the one cell stage; uninjected siblings were used as controls. **B.** Quantification of Pax7+ cells from the analysis shown in (**A**). The number of Pax7+ cells was significantly less in *hbegfa* crispants at 2dpi. Control and *hbegfa* crispant larvae showed comparable numbers of Pax7+ cells prior to injury. **C.** Survival curve. Tg(*actc1b*:NTR,2A,mCherry) larvae were treated with MTZ. *hbegfa^-/-^*larvae showed significantly reduced survival compared to sibling controls. **D.** Transverse sections of MTZ-treated Tg(*actc1b*:NTR,2A,mCherry) larvae at 4dpi, labelled for MyHC (green) and mCherry (red). *hbegfa^-/-^* larvae had significantly fewer myofibers than sibling controls. Data points represent average myofiber number per section from a single larva. **E.** Experimental design for systemic injection of human recombinant Hb-EGF. *pax7a^GFP,2A,NTR2.0^* larvae were injected through the duct of Cuvier at 4dpf and live imaged at 26hpi. Live images of *pax7a^GFP,2A,NTR2.0^*injected larvae show GFP+ cells in the trunk of Hb-EGF injected larvae. PBS injected siblings were used as controls. Arrows denote GFP+ cells. **F**. Quantification of injected larvae with Pax7 (GFP+) cells. n=20 per group. Error bars represent SEM. Significance was determined by unpaired students t-test (**B,D**) and Log-rank test (**C**). (ns) not significant, (*) p<0.01, (****) p<0.0001.

We recovered *hbegfa* alleles through the germ line that completely deleted the *hbegfa* coding region, and found *hbegfa^-/-^* larvae transgenic for Tg(*actc1b:*NTR,2A,mCherry) had reduced viability after systemic injury with a sharp decline in survival starting at 4dpi (Fig 6C). Histological analysis showed *hbegfa^-/-^* larvae had significantly fewer myofibers at 4dpi compared to controls (Fig. 6D). To understand if *hbegfa* is required for local muscle repair, we performed local needlestick on 4dpf larvae. Histological analysis of fast muscle showed local muscle repair was unaffected in *hbegfa^-/-^* larvae at 7dpi (Fig. S6B). Hb-EGF is thus required for systemic but not local muscle repair.

Our *hbegfa* crispant studies suggested Hb-EGF regulates MuSC proliferation (Fig. 6B). To functionally assess the role of Hb-EGF in MuSC activation, we injected human recombinant Hb-EGF protein into the systemic circulation of *pax7a^GFP,2A,NTR2.0^* larvae and assessed GFP expression 26hrs post injection (Figs. 6E, S6C). Larvae injected with Hb-EGF showed dose-dependent GFP expression, arguing Hb-EGF activates MuSC proliferation.

## Discussion

Our model of inducible systemic muscle injury revealed systemic and local injuries induce distinct molecular responses to direct muscle repair. Systemic muscle injury resulted in myofiber damage, which was predominantly repaired but not replaced with new myofibers. Transcriptomic analysis showed intercellular signaling strength was higher during systemic muscle repair relative to local repair. This differential signaling strength may reflect the need to orchestrate an organism-wide response to repair muscle that involves signaling across tissues. Hb-EGF expression was significantly enriched outside the myotome after systemic injury and was required for systemic repair. In addition, systemic delivery of Hb-EGF activated MuSC proliferation, arguing Hb-EGF can act over long distances to promote muscle repair. These studies demonstrate multiple regenerative programs are in place to ensure robust regeneration of injured skeletal muscle.

The systemic environment plays an important yet relatively unknown role in regulating MuSC function. Heterochronic parabiotic experiments, in which the circulatory systems of young and aged mice are mixed, revealed an incredible capacity for the young circulatory system to enhance MuSC proliferation and to reduce fibrogenic differentiation in aged mice(Hong et al., 2022). Conversely, the circulatory system from old mice increased fibrosis in young mice. These foundational studies revealed that the systemic environment provides an additional layer of regulation beyond the stem cell niche to control MuSC function. Although the circulating factors that direct age-related changes in MuSC function remain controversial (Hong et al., 2022), these studies demonstrate that MuSC activity can be remotely controlled over relatively long distances.

An essential clue that muscle injury induces an organism-wide response at the cellular level appeared in studies in which mammalian skeletal muscle was damaged in one limb, and MuSC activity was assessed in the uninjured, contralateral limb. Surprisingly, quiescent MuSCs in the contralateral limb transitioned into an “alert” state after injury, and were poised to activate and repair tissue damage quickly (Rodgers et al., 2014). At the molecular level, hepatocyte growth factor activator (HGFA) is induced by muscle injury, and activates human growth factor (HGF). HGF in turn acts at the stem cell niche where it directs MuSCs to transition into the alert state (Rodgers et al., 2014; Rodgers et al., 2017). HGFA treatment also appears to alert skin and hair follicle stem cell pools (Rodgers et al., 2017), arguing long-range signaling is a conserved strategy to enhance regeneration and repair.

In our model of systemic muscle repair, Hb-EGF acts over long distances to promote muscle regeneration. The primary tissue that expressed Hb-EGF in response to systemic muscle injury was the maturing epidermis and the MuSC population most affected in *hbegfa* crispants were tissue-resident MuSCs interstitially positioned among the fast myofibers. In addition, Hb-EGF protein injected into circulation induced MuSC proliferation at a distance far from the injection site. These data are consistent with a model in which long-range Hb-EGF signaling across tissues directs systemic muscle repair.

While the in vivo function of Hb-EGF during mammalian muscle regeneration has not been identified, Hb-EGF stimulated bovine MuSC proliferation in culture(Thornton et al., 2015). Outside the nervous system, Hb-EGF is produced by both macrophages and endothelial in the liver after ischemia injury and is required for hepatocyte proliferation during liver regeneration(Wu et al., 2024; Yang et al., 2024). In zebrafish, ependymal cells express Hb-EGF in response to spinal cord injury and delivery of recombinant Hb-EGF improved functional regeneration(Cigliola et al., 2023). Hb-EGF expression is also upregulated in response to olfactory epithelium injury, where it promotes proliferation of sensory neuron precursors and accelerates recovery(Sireci et al., 2024). Hb-EGF is thus a pro-regenerative factor under multiple injury contexts and activates the proliferation of reparative stem cells and cell-type specific progenitors. Although the molecular mechanisms by which injury activates Hb-EGF expression has been investigated in some detail(Cigliola et al., 2023), the intracellular pathways and targets that direct stem cell activation or progenitor proliferation in response to Hb-EGF are not known. Further investigation into these downstream pathways will be essential for understanding how MuSCs are activated in response to tissue cross talk and may reveal important insights into the mechanisms by which stem cells illicit specific responses to individual growth factors.

A most striking result from our study is that the repair of local muscle injuries did not require Hb-EGF, arguing Hb-EGF activates a MuSC response during systemic repair that is distinct from the response induced during local repair. Hb-EGF signaling may function at various levels to regulate the MuSC response to systemic injury. For example, Hb-EGF could act upstream of the local growth factors that activate MuSCs to amplify the magnitude of the overall MuSC response to enact systemic repair. On the other hand, Hb-EGF may signal directly to MuSCs to induce an intracellular response that alters the dynamics of proliferation and differentiation such that MuSCs undergo extended proliferation prior to differentiation. Increasing MuSC proliferation would ultimately provide more progenitors for systemic muscle repair. Our transcriptomic analysis argues Hb-EGF signals directly to MuSCs, but we cannot exclude a role for Hb-EGF in indirectly regulating MuSC activation. Since Hb-EGF was not required for local muscle repair, identifying the cellular and molecular targets of Hb-EGF signaling will provide an entry point for defining the regenerative programs that direct systemic repair.

If local and systemic muscle injuries induce distinct regenerative programs, then there must be a mechanism that evaluates the scale of the injury and elicits the appropriate response. Local injuries activate gene expression changes in remote tissues, suggesting injury information is delivered over long distances to coordinate an organism-wide response. For example, cardiac damage in zebrafish induces transcriptional changes in brain ependymal cells and kidney tubular cells through hormone signaling (Sun et al., 2022). The roles of long-range tissue interactions and inter-organ communication pathways are thus emerging as essential components of the regenerative response. It remains unclear if local and systemic muscle injuries activate inter-organ communication pathways that deliver injury information to distant tissues that in turn decipher the amplitude of an injury to activate the appropriate regenerative program. Our transcriptomic and genetic analyses however have revealed new insights into the organism-wide response to muscle injury and suggest that systemic repair requires multiple inputs outside of the MuSC niche.

While mammals can regenerate small-scale muscle injuries, large-scale muscle injuries such as trauma-induced muscle loss do not fully repair and can lead to debilitative disabilities (Sun et al., 2022). To improve large-scale muscle repair, bioengineering approaches are being developed for the localized delivery of bioactive molecules at the site of muscle loss (Haas et al., 2021). However, conflicting outcomes argue growth factor loaded scaffolds are not yet fully optimized to treat major muscle injuries (Gahlawat et al., 2024). Pro-regenerative molecules discovered in zebrafish can improve muscle outcomes following large-scale muscle injuries in mammals (Ratnayake et al., 2021). One exciting possibility is that the regenerative programs used for systemic muscle repair in teleosts are simply dormant in mammals. If this is the case, activators of systemic repair programs in zebrafish would be attractive candidates to induce analogous regenerative responses in mammals and treat large-scale muscle injuries.

## Acknowledgments

We thank Sharon Amacher for providing fish lines and helpful suggestions throughout the project and the Washington University School of Medicine Zebrafish Consortium for providing the facilities to generate and maintain the fish lines used in this study. A.N.J was funded by National Institutes of Health (R01AR070299), Washington University Musculoskeletal Research Center (NIH P30 AR074992, and a Pilot seed grant funding provided by the Washington University Center for Regenerative Medicine (CRM). M.H.M. was funded by National Institutes of Health (2R01NS113915 and 1R01NS123708).

## Author contributions

Conceptualization, H.C.D., V.M.S., M.H.M, A.N.J.; Data Curation, H.C.D., V.M.S., A.S., T.O. J.M., A.T.; Formal Analysis, H.C.D., V.M.S., A.S., T.O. J.M., A.T., M.H.M, A.N.J.; Funding Acquisition, M.H.M and A.N.J.; Investigation, H.C.D., V.M.S., A.S., T.O. J.M., A.T.; Methodology, H.C.D., V.M.S., A.S., J.M., A.T., M.H.M, A.N.J. Resources, M.H.M, A.N.J.; Writing Original Draft, A.T. and A.N.J.; Review & Editing, H.C.D., V.M.S., A.S., T.O. J.M., A.T., M.H.M, A.N.J.; Supervision, V.M.S, M.H.M and A.N.J.;

## Supplemental information

Table S1

Table S2

Table S3

## Methods

### Zebrafish

Zebrafish (*Danio rerio*) of the Tubingen genetic background were maintained at the Washington University Zebrafish Core Facility. All experimental protocols were approved by the Washington University School of Medicine Institutional Animal Care and Use Committee. Male and female larvae of the indicated ages were used for regeneration experiments. For all experiments, multiple cohorts (clutches) of injured fish were used. The number of animals for each experiment is reported in the figures or figure legends. The following published strains were used: Tg(- 80.0*myf5*:GFP) (Chen et al., 2007) and Tg(ubb:LOXP-RFP-STOP-LOXP-GFP) (Mosimann et al., 2011).

### Muscle injury

To induce a systemic injury, 50 larvae per experimental group were treated for 12 hours with 5mM of MTZ. Uninjured siblings were used as controls. Localized muscle injury was carried out as described (Gurevich et al., 2016). Zebrafish were anesthetized on a petri dish using 4 mg/mL tricaine solution (Syndel, 200-226). Localized injury was performed using size 0 insect pins (Roboz Surgical Instruments, RS-6081-35) at a 90° angle in three dorsal somites positioned above the cloaca. Injured fish were then transferred to a fresh petri dish with embryo water to recover.

### Generation of Transgenic Lines

#### Tg(*actc1b*:NTR,2A,mCherry)

actc1F and actc1R primers were used to amplify a 3.5kb region proximal to the *actc1b* promoter. This enhancer was previously characterized as skeletal muscle specific(Higashijima et al., 1997). The genomic fragment was cloned into pCR2.1, then subcloned into a BamHI digested PCS2-mCherry-Nitroreductase (NTR) plasmid (Zhou et al., 2023) to generate pCS2.*actc1b*:NTR.2A.mCherry. This plasmid was co-injected with I-SceI into zebrafish embryos at the one-cell stage. Founders were identified by mCherry expression and confirmed by genotyping using actc1genoF and NTRgenoR primers. Ten founder lines were tested for muscle injury and two of these founders were maintained for further studies.

#### Tg(*actc1b*:NTR,2A)

The pCS2.*actc1b*:NTR.2A.mCherry plasmid was digested with XhoI to remove the mCherry coding sequence and ligated. The plasmid was injected as described above. Ten founder lines were tested for muscle injury and two of these founders were maintained for further studies.

#### Tg(*actc1b*:CreER^T2^)

Two EcoR1 sites were added to the 3.5kb *actc1b* enhancer using standard PCR, cloned into pCR2.1., and then subcloned into EcoR1 digested pCS2.Flag.CreER^T2^ (Zhou et al., 2023) to generate pCS2.*actc1b*:CreER^T2^. The plasmid was co-injected with I-SceI into zebrafish embryos at the one-cell stage. Founders were genotyped with CreF and CreR primers. Twenty founder lines were crossed with ubb:LOXP-RFP-STOP-LOXP-GFP and offspring were treated with 5uM tamoxifen (Sigma T5648) for 16hr to identify two lines with robust recombination activity. Neither line activated GFP in the absence of tamoxifen.

Double and triple transgenic lines were generated using standard genetic crosses.

### CRISPR/Cas9 mutagenesis

CRISPR/Cas9 design and mutagenesis was followed according to(Hoshijima et al., 2019; Klatt Shaw et al., 2021). crRNAs were designed using CHOPCHOP(Labun et al., 2019) against the GRCz11 zebrafish genome. crRNA sequences with no off-target sites, minimum mismatches and highest efficiency were chosen. Alt-R modified crRNAs ordered from IDT (2 nmol scale) and tracrRNAs (IDT, 1073190) were resuspended according to manufacturer’s specifications. tracrRNA and crRNA were mixed and diluted to a final concentration of 50μM, annealed by heating to 95°C and gradual cooling at 2°C/second to 25°C, and stored at −20°C. For injections, duplexed dgRNAs were diluted 1:1 in duplex buffer (IDT, 1072570) to a working concentration of 25 μM. Equal volumes of dgRNA and Cas9 (IDT, 1081058) were incubated with 37.5mM KCl and injection dye for 37°C for 5 minutes. For co-injections with two guides, Cas9 was added in equal molar concentration as the total gRNA concentration. Embryos at the one-cell stage were injected with ∼1nL of CRISPR/Cas9 solution.

#### _pax7a_^GFP,2A,NTR2.0^ _knock-in_

A pCS2.pax7a.GFP-2A-NTR2.0 construct was generated by HiFi Assembly (New England Biolabs, E5520S) comprised of two 1kb genomic homology arms that flanked the *pax7a* stop codon and the GFP-2A-NTR2.0 cassette (Sharrock et al., 2022). To generate the GFP-2A-NTR2.0 knock-in, the crRNA *pax7a*.KI was duplexed and mixed with Cas9, KCl, and injection dye as described. The template plasmid pCS2.pax7a.GFP-2A-NTR2.0 (25ng/μl final concentration) and non-homologous end joining inhibitor NU7441 (Selleckchem, 50mM final concentration) were then added to the injection mixture. Embryos at the one-cell stage were injected with ∼1nL of CRISPR/Cas9 solution. Founders were identified by PCR screening F_0_ larvae for GFP (using GFPF and GFPR primers) and then confirmed with junctional primers that spanned the *pax7a* locus and the GFP-2A-NTR2.0 cassette (using Pax7AF and Pax7AR primers).

#### *smyhc1^sfgfp^*knock-in

A pMK-RQ.smyhc1.GFP construct (generated by Invitrogen) comprised of two 1kb genomic homology arms that flanked the *smyhc1* stop codon and a 714bp superfolder GFP coding sequence was used as a template for homology dependent repair. To generate the GFP knock-in, the crRNA *smyhc1*.GFP.KI was injected with the template plasmid pMK-RQ.smyhc1.GFP and the injection cocktail described above.

#### hbegfa deletion

hbegfa.1 and hbegfa.2 crRNAs targeting the first and last coding exons of *hbegfa* were duplexed and injected as previously described. hbegfadelF, hbegfadelR, and hbegfahetR primers were used to genotype the F_1_ and identify founders. Two founder lines were maintained.

### Recombinant protein injection

4dpf larvae were anesthetized with tricaine and positioned laterally in custom molds. Injection needles were filled with PBS or Hb-EGF protein (Sigma E4643) and 1μl Dextran-Alexa 594 (Invitrogen D22913) and larvae were injected in the duct of Cuvier as described (Benard et al., 2012). Larvae were live imaged 24hr after injection.

### Live imaging

Larvae were immobilized in imaging chambers with 1% low melt agarose. Fish water supplemented with tricaine was added to the chamber after the agarose set. Live images were taken using a Zeiss LSM 800 confocal microscope or a Nikon W1 CSU SoRa spinning disk confocal.

### Histology

Larvae were either dissected to remove heads for genotyping or fixed whole with 4% para-formaldehyde in PBS for 1hr. Samples were then rinsed 4X for 10min in PBT, and then transferred to 30% sucrose in PBS and incubated overnight at 4 °C. Samples were embedded in tissue freezing medium (General Data Healthcare. Cat# TFM-C), frozen on dry ice, and stored at −80 °C. 16 µm cross cryosections were generated for downstream application. Tissue sections were imaged using a Zeiss LSM 800 confocal microscope for immunofluorescence and HCR.

For Hybridization chain reaction fluorescent in-situ hybridization (HCR-FISH), probes were designed against target mRNAs using an automated program(Kuehn et al., 2022), and ordered from IDT as pooled oligonucleotides (50pmol scale). The HCR protocol was adapted from(Choi et al., 2018; Shaw et al., 2024). Slides were dried on a warmer for an hour before dehydrating for 3 minutes each in 50% ethanol in PBT, 95% ethanol in PBT and 100% ethanol. They were hydrated in 50% PBT in ethanol, 95% PBT in ethanol and 100% PBT. Slides were treated with 0.2% TritonX-100 in PBS, placed in PBT five times for 5 minutes. After drying the slides, samples were enclosed within a hydrophobic barrier. For blocking, slides were incubated for 1 hour at 37 °C in 200 μL of pre-warmed hybridization buffer (Molecular Instruments) per slide. Probes were prepared by adding 1μL of each desired probe with 200μL of pre-warmed hybridization buffer. The pre-hybridization buffer was replaced with the probe in hybridization buffer and sections were incubated at 37°C in dark for 48hrs. Slides were washed in 100μL of wash buffer (Molecular Instruments) at room temperature four times for 5min followed by 5X SSCT (3M NaCl, 0.3 M Sodium Citrate, 0.1% Tween-20, pH 7.0) twice for 5min. The sections were incubated in 200μL of amplification buffer (Molecular Instruments) per slide for 1hr at room temperature. Prior to amplification, hairpin RNAs were preheated for 90 seconds at 95°C and cooled for 30min. For amplification, each hairpin RNA was diluted 1:50 in amplification buffer and incubated at room temperature in dark for ∼24hrs. Samples were washed twice for 5min in 5X SSCT followed by 20X SSC at room temperature. Sections were mounted in 60 μL Fluoromount-G, incubated at 4°C overnight and sealed with nail polish.

Immunohistochemstry (IHC) was carried out as described in The Zebrafish Book (Westerfield, 2000). Briefly, tissue samples were restricted within a hydrophobic boundary with an ImmEdge barrier pen (Vector Laboratories. Cat# H-4000) and rehydrated in PBT (0.1% Tween-20 in PBS). After two 5-min washes, samples were treated with a mix of 0.1% Triton in PBS and rinsed in PBT four times for 5min. Tissue sections were blocked in 5% goat serum, 0.1% Triton in PBS for an hour at room temperature. Samples were incubated overnight with primary antibody diluted in 5% goat serum in PBT at 4°C in the dark followed by four 5min washes with PBT. Samples were then incubated in secondary antibody in 5% goat serum in PBT for 2 hours at room temperature in the dark followed by four 5min washes with PBT. Samples were incubated with 1 μg/mL of Hoechst for 5 minutes and rinsed twice with PBT for 5 minutes.

Primary antibodies used in the study were: mouse anti-4D9 (DSHB, 1:10), mouse anti-PAX7 (DSHB, 1:10), mouse anti-MF 20 (DSHB, 1:400), rabbit anti-DsRed (Living Colors, 632496, 1:400), chicken anti-GFP (Aves Labs, cat# GFP-1020, 1:1500), rabbit anti-D175 (Cell Signaling Technology, 966, 1:500). Secondary antibodies used were Alexa Fluor 488 goat anti-mouse (Invitrogen, # A-21121, 1:250), Alexa Fluor 594 goat anti-rabbit (Invitrogen, A-11072, 1:250), Alexa Fluor 488 goat anti-chicken (Invitrogen, cat# A-11039; RRID: AB_142924, 1:250)

### Quantification

IHC and HCR-FISH was performed on 3-5 serial transverse sections per fish. Quantification was performed on single-plane projected confocal images. The cell counter tool in ImageJ was used to quantify the number of cells that expressed specific proteins. To quantify HCR fluorescence, user-defined color thresholds were set manually in ImageJ. Integrated density was quantified for fluorescence signals inside a region of interest using the “Analyze Particles” command.

### Statistical Analyses

Statistical analyses for histology and functional assays were performed with GraphPad Prism software package and included unpaired student’s t-test (with Welch’s correction in cases of non-Gaussian distribution) and one-way ANOVA. Appropriate corrections for multiple pairwise comparisons among groups were applied.

### Isolation of single cells from larval zebrafish

15 larvae per cohort were dissected to remove the head, caudal fin, and internal organs and then processed as described (Farnsworth DB 459 2020). For tissue lysis, 500μL of dissociation buffer (2ug/ul Collagenase P, 2mM CaCl2, 0.2 ul DNaseI, 0.25% Trypsin, in 1X PBS) was added to the samples and incubated at 28 °C until samples were visually dissociated (∼15min). Dissociation was stopped with equal volume of stop buffer (10% FBS, 0.001mM EDTA, in 1X PBS). Samples were centrifuged at 750rcf for 7min at 4°C, the supernatant was discarded, and the pellet was resuspended in 100μL of resuspension buffer (1% FBS, 2mM CaCl2, 1X Penicillin/Streptomycin, in DMEM). The suspension was strained through 20μm cell strainer (Pluriselect USA, 43-10020-40), centrifuged once (500rcf, 1 min, 4°C) to pass cells through the strainer, and centrifuged a second time (750rcf, 7 min, 4°C) to pellet the cells. The cells were resuspended in 100μL of resuspension buffer, centrifugated (750rcf, 7min, 4°C), and the pellet was resuspended in 50μL 0.04% BSA in PBS. 5μl of cells were then combined with 5μl of Hoechst (Invitrogen, H3570), incubated for 10min, and transferred to a hemocytometer (Bulldogbio, NC1731934) to determine cell concentration. An additional 5μl of cells were incubated with Trypan blue (Sigma, T8154) for 5min, transferred to a hemocytometer, and counted to assess cell viability. Samples with > 70% viability were then submitted for single cell sequencing.

### Single cell RNA-sequencing

For scRNA-seq, 30 µl of resuspension solution containing isolated cells at a concentration of ∼1000 cells/µl were submitted to the Genome Technology Access Center in the McDonnell Genome Institute at Washington University. One biological replicate was used for each time point. cDNA was prepared after GEM generation and barcoding, followed by the GEM-RT reaction and bead cleanup steps. cDNA was amplified for 11−13 cycles then purified using SPRIselect beads. Purified cDNA samples were then run on a Bioanalyzer to determine cDNA concentration. GEX libraries were prepared as recommended by the 10x Genomics Chromium Single Cell 3’ Reagent Kits User Guide (v3.1 Chemistry Dual Index) with appropriate modifications to the PCR cycles based on the calculated cDNA concentration. For sample preparation on the 10x Genomics platform, the Chromium Next GEM Single Cell 3’ Kit v3.1, 16 rxns (PN-1000268), Chromium Next GEM Chip G Single Cell Kit, 48 rxns (PN-1000120), and Dual Index Kit TT Set A, 96 rxns (PN-1000215) were used. The concentration of each library was accurately determined through qPCR utilizing the KAPA library Quantification Kit according to the manufacturer’s protocol (KAPA Biosystems/Roche) to produce cluster counts appropriate for the Illumina NovaSeq6000 instrument. Normalized libraries were sequenced on a NovaSeq6000 S4 Flow Cell using the XP workflow and a 50 × 10 × 16 × 150 sequencing recipe according to manufacturer protocol. A median sequencing depth of 50,000 reads/cell was targeted for each Gene Expression Library.

### Aligning scRNA-seq reads

After sequencing, the Illumina output was processed using the CellRanger (v8.0.1) pipeline to generate gene-barcode count matrices. A custom reference genome was made with the “cellranger mkref” command, using the fasta file of zebrafish reference genome GRCz11 constructed from the Ensemble genome build (https://useast.ensembl.org/Danio_rerio/Info/Index) and the sorted Gene Transfer Format file (v4.3.2) from the improved zebrafish transcriptome annotation (Lawson et al., 2020). Base call files for each sample from Illumina were demultiplexed into FASTQ reads. Then, the “cellranger count” pipeline was used to align sequencing reads in FASTQ files to the custom reference genome. Both exon and intron sequences were included during the alignment. The filtered gene-barcode count matrices generated by “cellranger count” was used for downstream analysis.

### Quality control

DecontX package (v1.2.0) was used to remove droplets containing aberrant counts of ambient mRNA (Yang et al., 2020). The function “decontx” was used on the raw “RNA” counts to obtain “decontX” count data, following the default pipeline. DecontX counts were used as default for preprocessing and normalization. After ambient RNA removal, DoubletFinder (v2.0.4) was used to identify doublets formed from transcriptionally distinct cells (McGinnis et al., 2019). Optimal pK values were calculated from the outputs of function “paramsweep”. Then, “doubletFinder” function was used to predict doublet cells, where 50 principal components and pN value of 0.25 were given as input along with the previously calculated pK value (30). All the cells predicted as doublets were removed.

### Integrated analysis of scRNA-seq dataset

Datasets were integrated and analyzed using Seurat (v4.4.0) package with R (v4.4.2)(Stuart et al., 2019; Team, 2024). Each sample count matrix was filtered for genes that were expressed in at least 3 cells and cells expressing at least 200 genes, followed by cell quality assessment using commonly used QC matrixes (Ilicic et al., 2016). Cells having a unique number of genes between 200 and 6500, mitochondrial gene percentage <5 and total number of reads (UMIs) between 300 and 35000 were used for downstream processing. Each dataset was independently normalized and scaled using the “SCTransform” function, which is an improved method for normalization, that performs a variance-stabilizing transformation using negative binomial regression (Hafemeister and Satija, 2019). Standard integration workflow of Seurat was used to identify shared sources of variation across experiments as well as mutual nearest neighbors (Butler et al., 2018; Haghverdi et al., 2018). Integration features were selected based on the top 6000 highly variable features using “SelectIntegrationFeatures” function (nfeatures = 6000), which was used as input for the “anchor.features” argument of the “FindIntegrationAnchors” function. PCA analysis was performed on the 6000 variable features and the top 50 principal components selected based on the elbow plot heuristic, which measures the contribution of variation in each component. These 50 principal components were used in “FindNeighbors” and “FindClusters” functions to perform graph-based clustering on a shared nearest neighbor graph (Levine et al., 2015; Xu and Su, 2015). Louvain algorithm was used for modularity optimization in cell clustering using “FindClusters” function. The resolution parameter (res = 0.3) that determines the granularity of clustering was selected by visually inspecting clusters with resolutions ranging between 0.1 and 2.0 as well as clustree graph (Zappia and Oshlack, 2018). Uniform Manifold Approximation and Reduction (UMAP) was used for non-linear dimensional reduction of the first 50 principal components and visualize the data using “RunUMAP” function (Becht et al., 2018). Data was graphed using different plot functions, such as “DimPlot”, “VlnPlot”, “FeaturePlot”, “Dotplot” and “DoHeatmap”, to view the cell cluster identity and marker gene expression. Cell proportions were extracted using the “table” and “prop.table” functions. Differential gene expression for individual clusters was identified using Wilcoxon rank sum test in the “FindAllMarkers” function. Marker genes detected in at least 25% of the clustered cells and had a positive average log_2_(FC) were reported.

### Cluster identification using differentially expressed markers

We used a zebrafish single atlas (Sur et al., 2023) to generate a database of markers for each tissue and cell type (Table S1). The differentially expressed markers of each cluster were crossreferenced with our compiled database using our “DE-Marker-Scoring” algorithm (Saraswathy et al., 2024). For every matching marker gene, one point was given to the respective cluster under the column name with matching cell identity. Iteration over every marker gene was performed to generate a scoring matrix with varying points for each cluster against the different cell identities compiled in the database (Table S2, sheet: scoring). The “phyper” function in R was then used to calculate one-tailed binomial probabilities using hypergeometric distribution for the total score obtained from each cluster against each cell identity in the database (Table S2, sheet: Binomial probability). −log10 of probability values were obtained for plotting the heatmap (Table S2, sheet:−log10P). The resulting values were scaled from 0 to 100 and plotted as a heatmap using GraphPad prism. Each cluster was given an identity based on the maximum −log10 p-score obtained in the heatmap. The top DE markers of clusters with ambiguous scores were manually annotated using the “ident genes” from the zebrafish single atlas (Sur et al., 2023) and literature search (McKellar et al., 2021). Top DE markers generated for each cluster is given in (Table S3, sheet:topDEmarkers.all.clusters) “RenameIdents” function was used to assign identity to each cluster. To confirm the assigned cluster identities, enrichment of classical markers of respective cell types were tested using Dot plot.

### Subset analysis of muscle clusters

Muscle cells identified from the complete dataset were subclustered using the “subset” function for subcluster analysis. The muscle subset was again normalized and scaled using the “SCTransform” function with glmGamPoi method (Ahlmann-Eltze and Huber, 2021). Fifty principal components were used and the resolution parameter was set to 0.4. Downstream analysis was done as described above for the integrated analysis. The top DE markers generated for each muscle subcluster using “FindAllMarkers” function is given in (Table S3,sheet: topDEmarkers.muscle.clusters).

### Cell-cell interaction assay

The R package CellChat (v2.1.2) was used to evaluate regenerative cell-cell interactions after local and systemic muscle injury (Jin et al., 2021). CellChat models the probability of cell-cell communication by integrating our gene expression data with a database of known interaction between signaling ligands, receptors, and their cofactors (CellChatDB). The RNA data was used to create CellChat object using “createCellChat” function, followed by the recommended preprocessing functions with default parameters for the analysis of individual datasets. Truncated mean method with 10% trimmed observation was used to compute average gene expression per cell group for the complete dataset. The default trimean method was used to compute average gene expression for the CellChat analysis. CellChatDB.zebrafish was used to infer cell-cell communication. All categories of ligand-receptor interactions in the database were used in the analysis. Communications involving less than 10 cells were excluded. The “netAnalysis_computeCentrality” function was used to calculate network centrality scores at each time point. Functions such as “netVisual_circle”, “netAnalysis_contribution”, “netVisual_aggregate”, “netVisual_bubble”, “netAnalysis_signalingRole_heatmap”, and “netAnalysis_signalingRole_scatter” were used to generate different plots used in this paper.

Primers used in this study were:

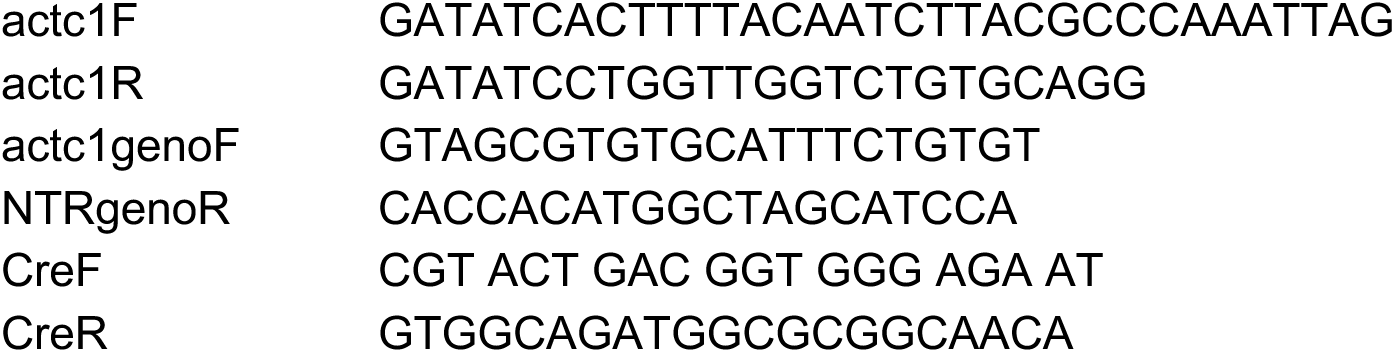
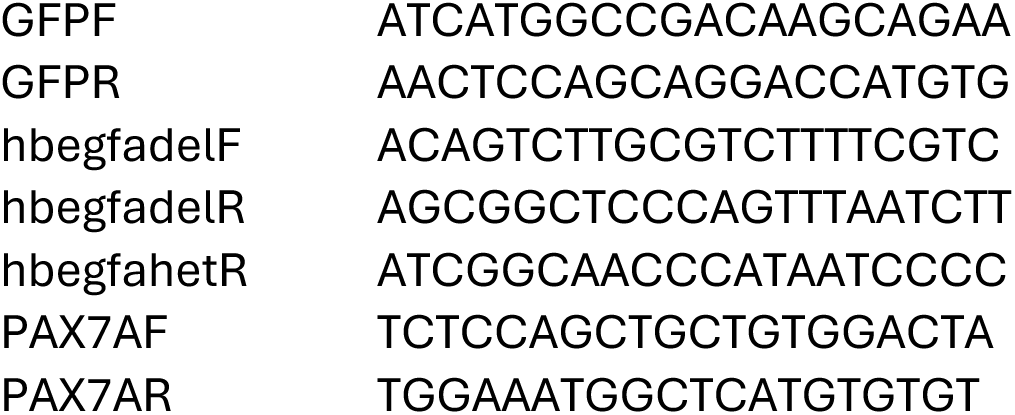

CRISPR (crRNA) guides used in this study were:

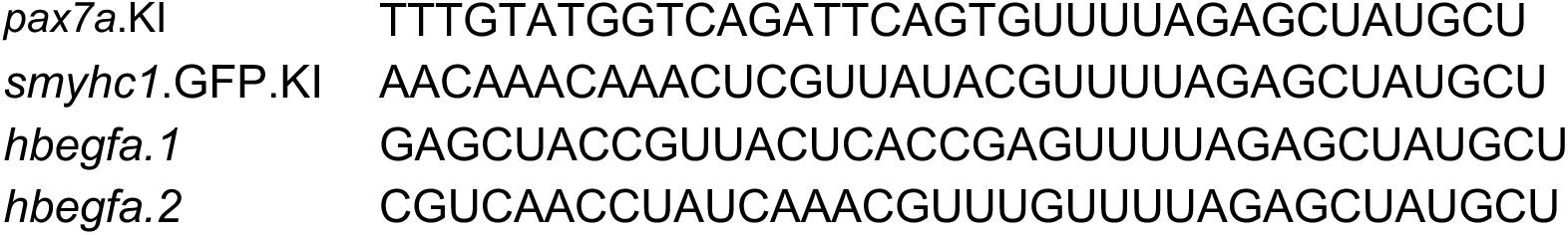

---

**Figure S1.**
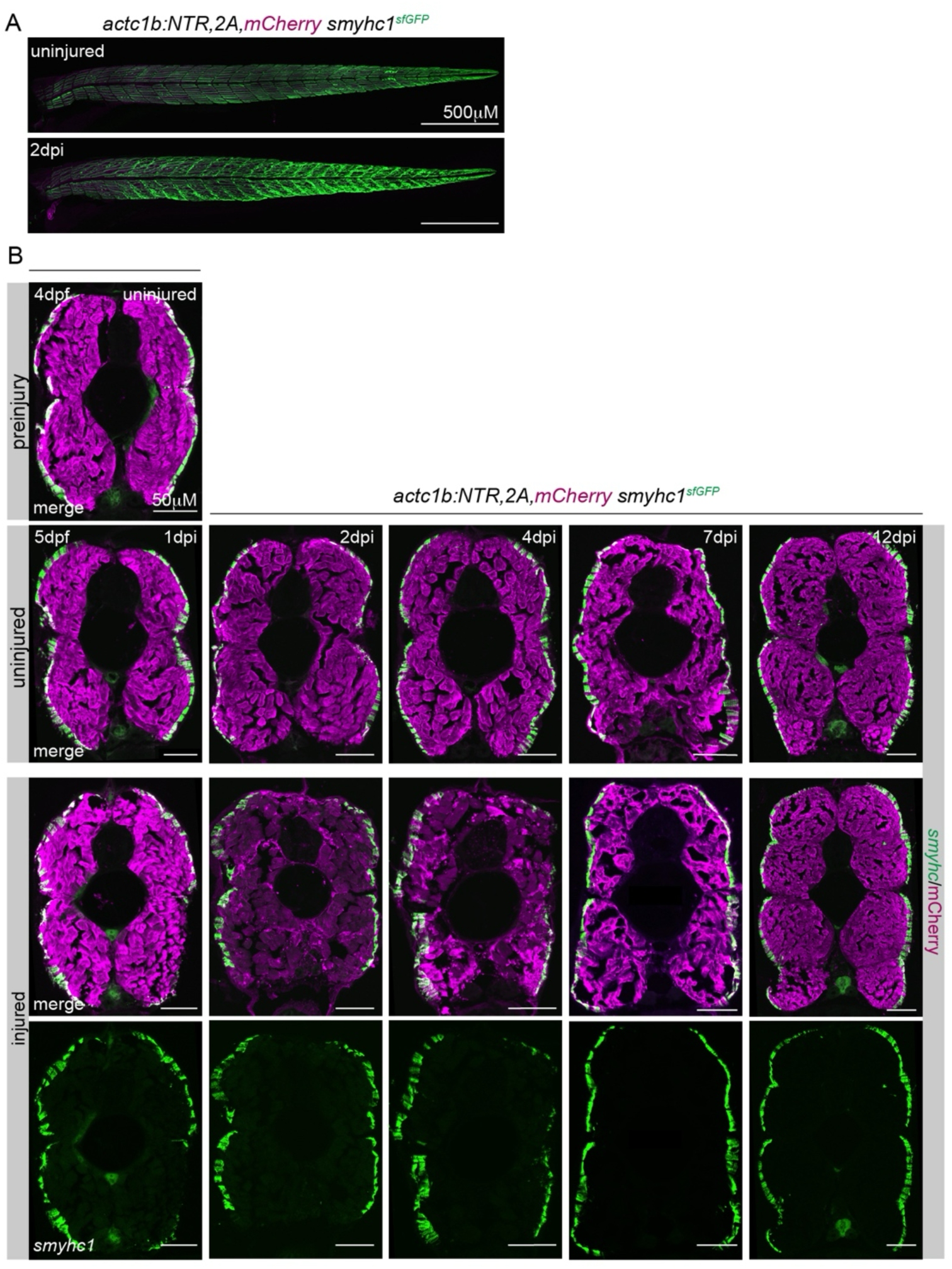
*smyhc1* is expressed in slow but not fast myofibers during systemic repair. **A.** Live images of *smyhc1^sfgfp^* Tg(*actc1b*:NTR,2A,mCherry) larvae treated with MTZ at 4dpf for 12hr. The pattern of Smyhc1:GFP (green) was disrupted in injured larvae at 2dpi. **C.** Regeneration time course. Transverse sections of *smyhc1^sfgfp^* Tg(*actc1b*:NTR,2A,mCherry) larvae from pre-injury and regeneration timepoints labelled for mCherry (magenta) and Smyhc1.GFP (green). Smyhc1.GFP was maintained in slow myofibers throughout systemic repair and was not activated in fast myofibers.

**Figure S2.**
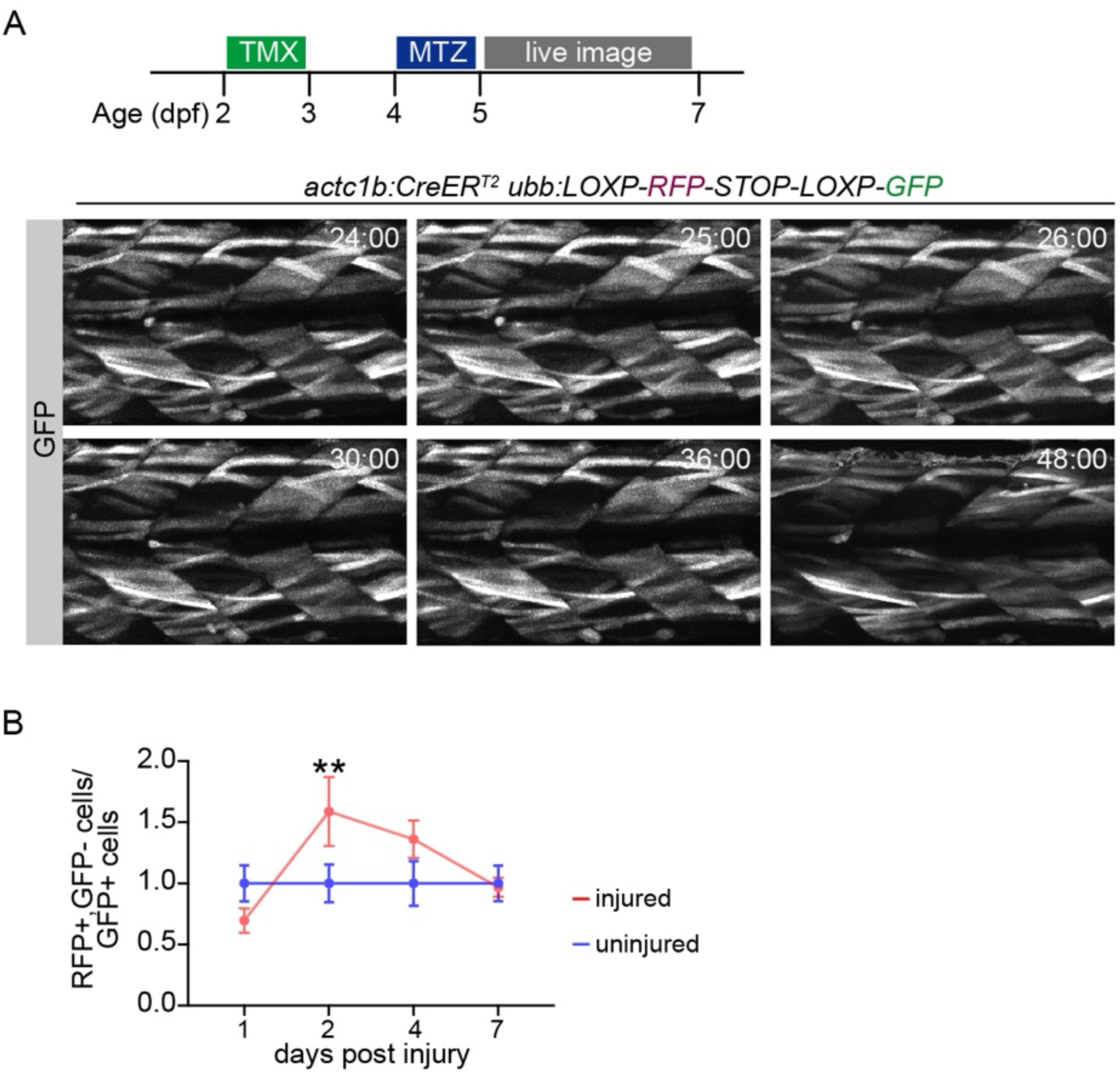
Injured myofibers survive systemic injury. **A.** Larvae triple transgenic for Tg (*actc1b*:NTR), (*actc1b*:CreER^T2^),(*ubb*:LOXP-RFP-STOP-LOXP-GFP) were treated with TMX for 24hr at 2dpf, and then treated with MTZ for 24hr at 4dpf. GFP+ larvae were live imaged for 24hr (n=3). Labelled myofibers survived systemic injury. **B.** Quantification of GFP+ and GFP-myofibers from lineage traced larvae collected at 2-, 4- and 7dpi. While a majority of myofibers survive systemic injury, there was a significant increase in GFP-myofibers at 2dpi suggesting some *de novo* myofiber generation contributes to muscle regeneration. n≥3 per treatment per timepoint.

**Figure S3.**
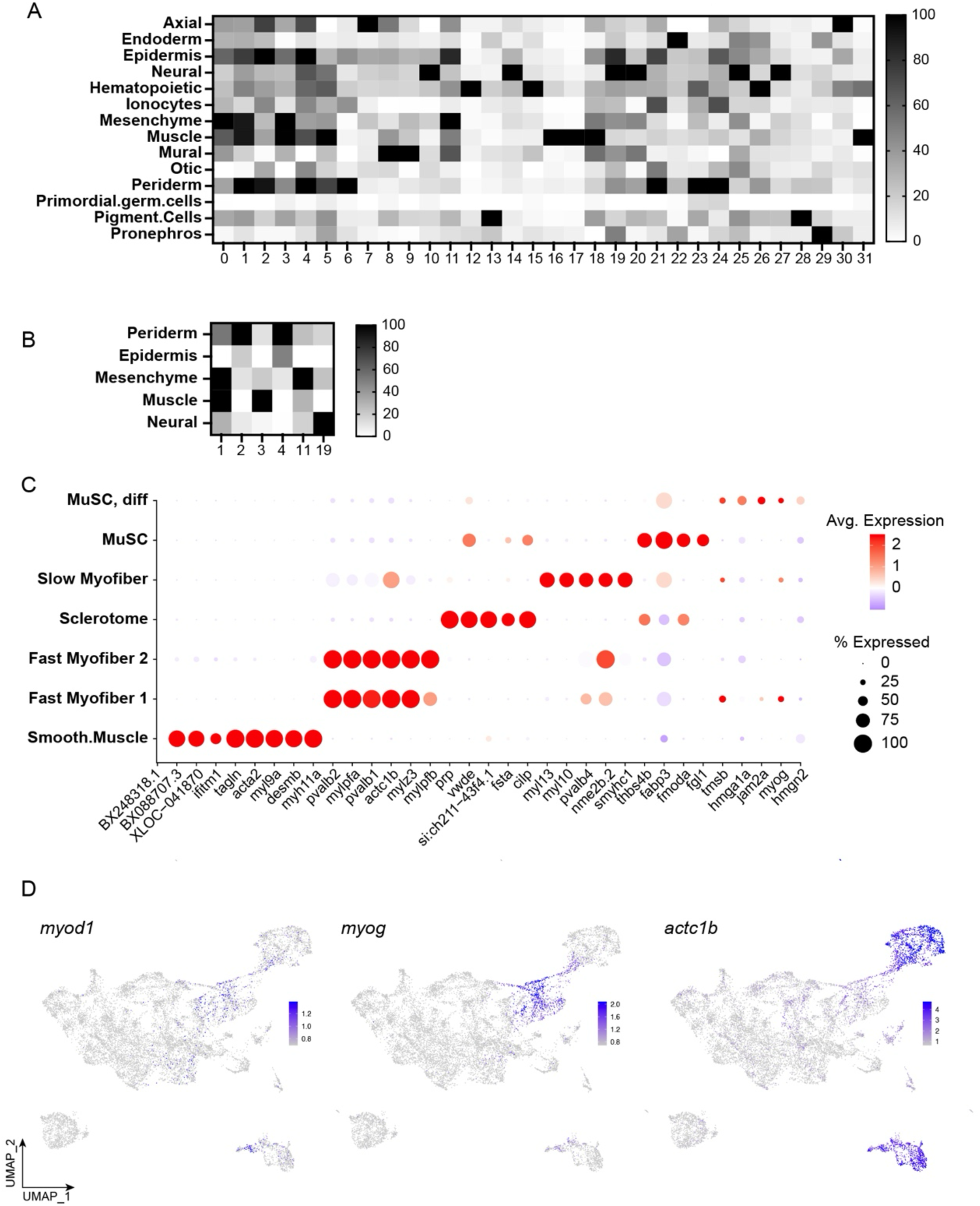
Cluster identification and validation. **A.** Heatmap to define each cluster by tissue type using the DE-Marker-Scoring” algorithm. **B.** Three ambiguous clusters were manually annotated using the “ident genes” from the zebrafish single atlas (Sur et al., 2023). **C.** Marker gene expression of muscle cell types in the complete data set. Dot plot shows top markers enriched in each cell type. Dot colors and diameters represent average gene expression and the percent of cells expressing a given marker. **D.** Feature plots showing the distributions of differentiating myoblast markers (*myod*, *myog*) and the terminal differentiation marker *actc1b*.

**Figure S4.**
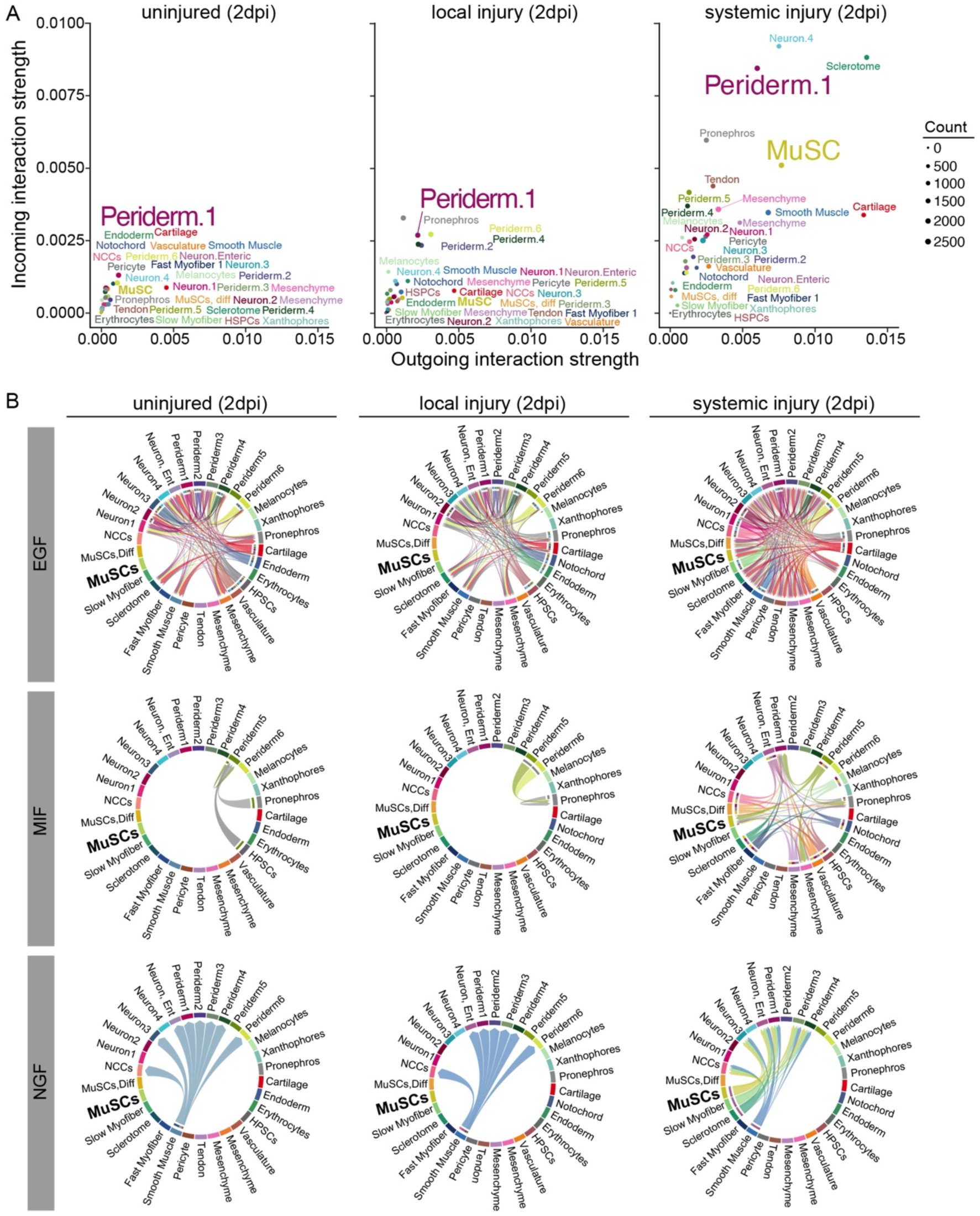
Systemic injury activates unique cell-cell communication networks. **A.** Relative incoming and outgoing interaction strengths by cluster in uninjured, locally injured, and systemically injured samples at 2dpi. More clusters showed strong interactions in response to systemic injury than local injury or no injury. **B.** Chord diagrams showing Epidermal growth factor (EGF), Macrophage migration inhibitory factor (MIF), and Nerve growth Factor (NGF) intercellular communication at 2dpi. Notice only systemic injury induced EGF and MIF signaling to MuSCs or NGF signaling out of MuSCs.

**Figure S5.**
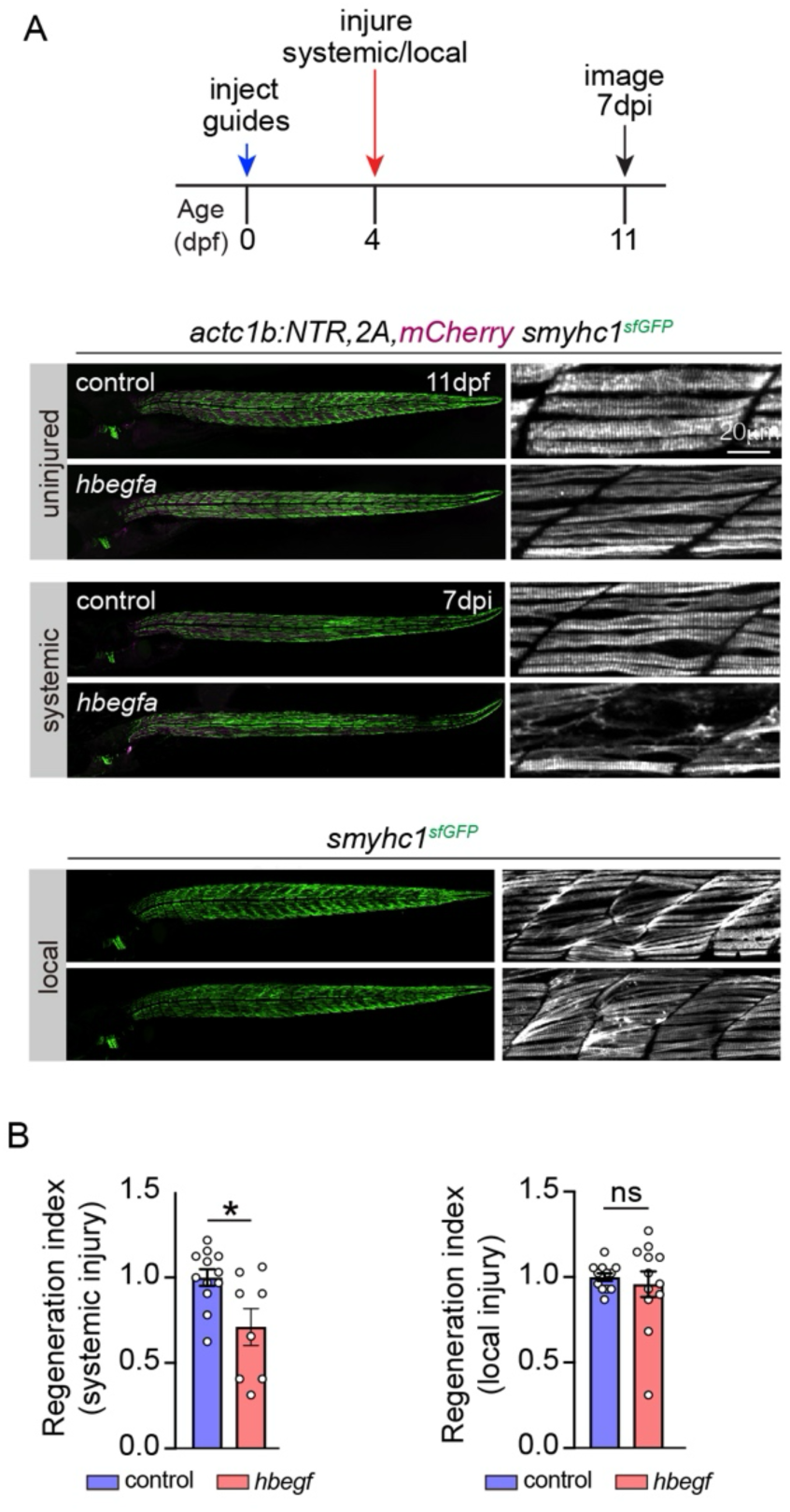
*hbega* is dispensable for local repair. **A.** Experimental design for CRISPR-mediated deletion of *hbegfa*. Injected embryos were injured at 4dpf and collected at 7dpi for live imaging of Smyhc1:GFP (green). *hbegfa* crispants were defective in slow myofiber repair after systemic but not local injury. Uninjected siblings were used as controls. Uninjured crispants and control larvae showed similar muscle morphology (top panel). **B.** Quantification of regeneration. The number of Smyhc1:GFP positive myofibers were counted in three somites per larvae at 7dpi to calculate a regeneration index. *hbegfa* crispants showed significantly less slow myofiber repair after systemic injury compared to controls. Slow myofiber repair in response to local injury was comparable between crispants and controls. Data points represent individual larva. Error bars represent SEM. Unpaired students t-test. (ns) not significant (*) p<0.05.

**Figure S6.**
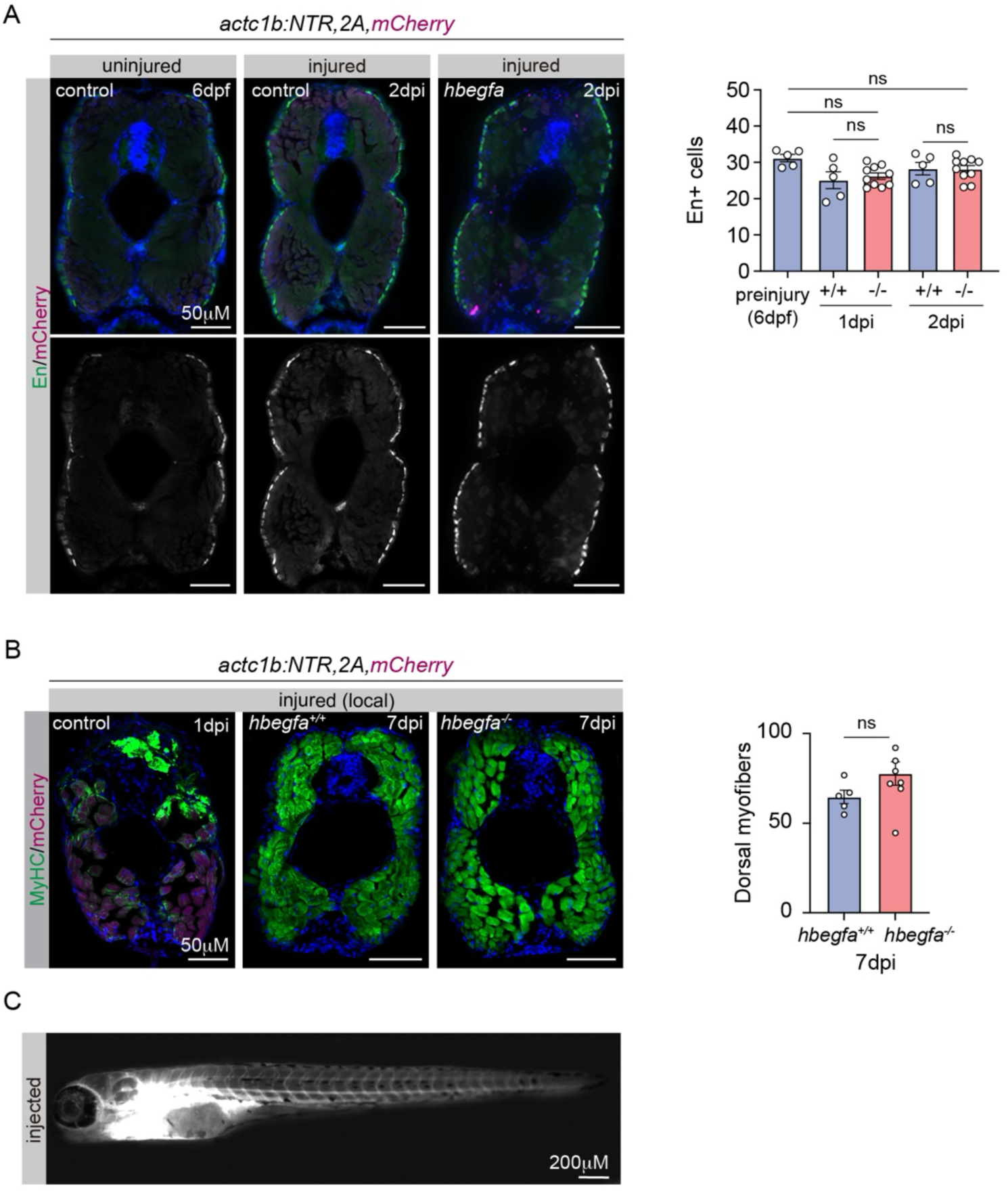
Hb-EGF does not En1+ MuSCs or local muscle repair. **A.** Transverse sections of the trunk musculature of Tg(*actc1b*:NTR,2A,mCherry) larvae labelled for En1 (green) at 2dpi. The number of En1 expressing cells in *hbegfa* crispants was comparable to uninjected sibling controls at 1- and 2-dpi. **B.** Transverse sections of Tg(*actc1b*:NTR,2A,mCherry) larvae at the site of local (needlestick) injury at 1- and 7-dpi, labelled for MyHC (green) and mCherry (red). Local injury caused MyHC aggregation at the injury site one day after injury. *hbegf^+/+^* and *hbegfa^-/-^* larvae showed comparable myofiber repair at the site of injury (dorsal myotome) by 7dpi. Data points represent average myofiber number per section from a single larva. **C.** Live images of *pax7a^GFP,2A,NTR2.0^* injected with fluorescent Dextran-594 through the duct of Cuvier at 4dpf and live imaged 20 minutes later. Injected larvae showed 594-fluorescence throughout the vasculature. Error bars represent SEM. Significance was determined by unpaired students t-test. (ns) not significant.

